# Lipid metabolism drives allele-specific early-stage hypertrophic cardiomyopathy

**DOI:** 10.1101/2023.11.10.564562

**Authors:** Arpana Vaniya, Anja Karlstaedt, Damla Ates Gulkok, Tilo Thottakara, Yamin Liu, Sili Fan, Hannah Eades, Ryuya Fukunaga, Hilary J. Vernon, Oliver Fiehn, M. Roselle Abraham

## Abstract

Hypertrophic cardiomyopathy (HCM) results from pathogenic variants in sarcomeric protein genes, that increase myocyte energy demand and lead to cardiac hypertrophy. But it is unknown whether a common metabolic trait underlies the cardiac phenotype at early disease stage. This study characterized two HCM mouse models (R92W-TnT, R403Q-MyHC) that demonstrate differences in mitochondrial function at early disease stage. Using a combination of cardiac phenotyping, transcriptomics, mass spectrometry-based metabolomics and computational modeling, we discovered allele-specific differences in cardiac structure/function and metabolic changes. TnT-mutant hearts had impaired energy substrate metabolism and increased phospholipid remodeling compared to MyHC-mutants. TnT-mutants showed increased incorporation of saturated fatty acid residues into ceramides, cardiolipin, and increased lipid peroxidation, that could underlie allele-specific differences in mitochondrial function and cardiomyopathy.

**Graphical Abstract:** 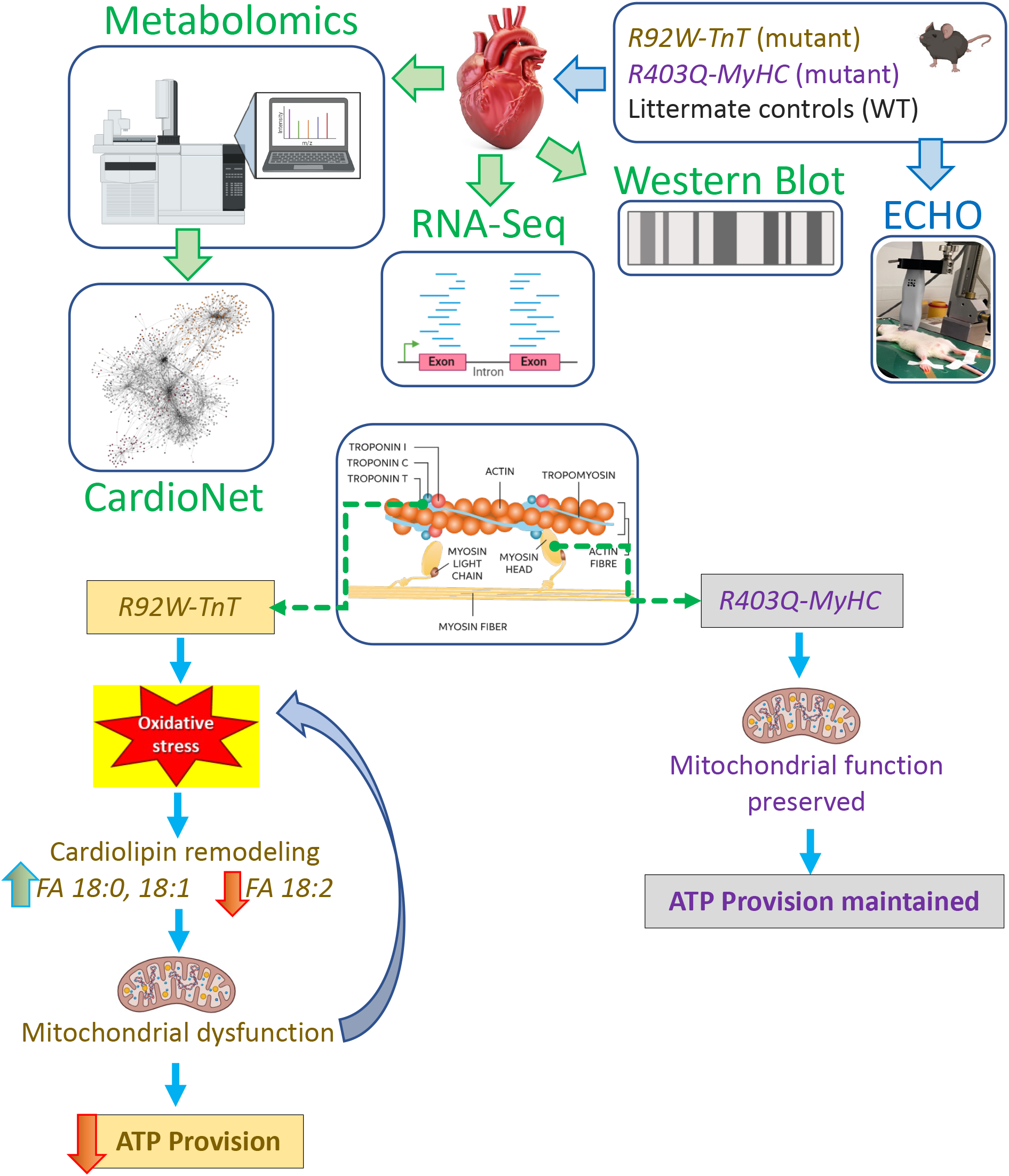
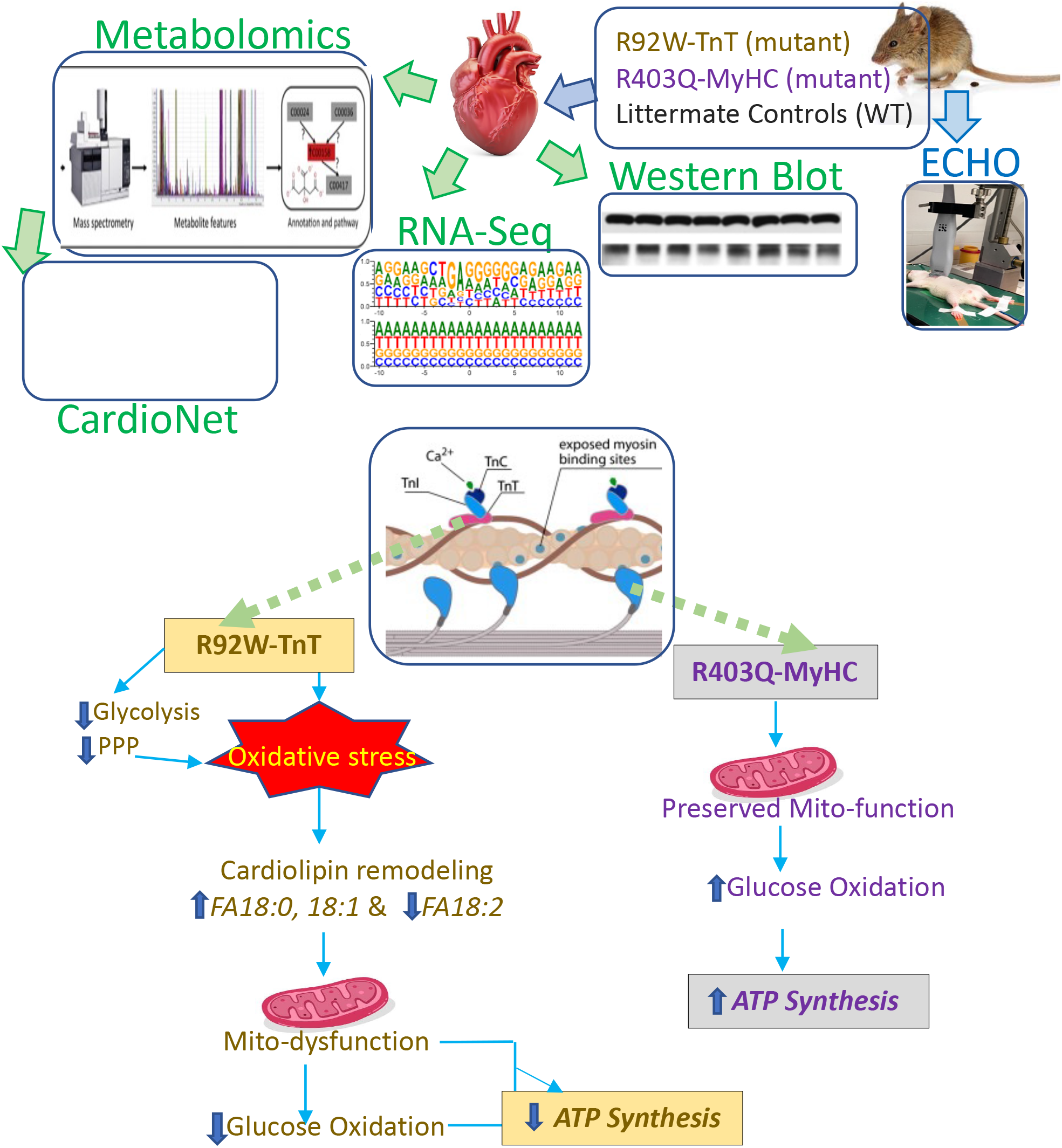
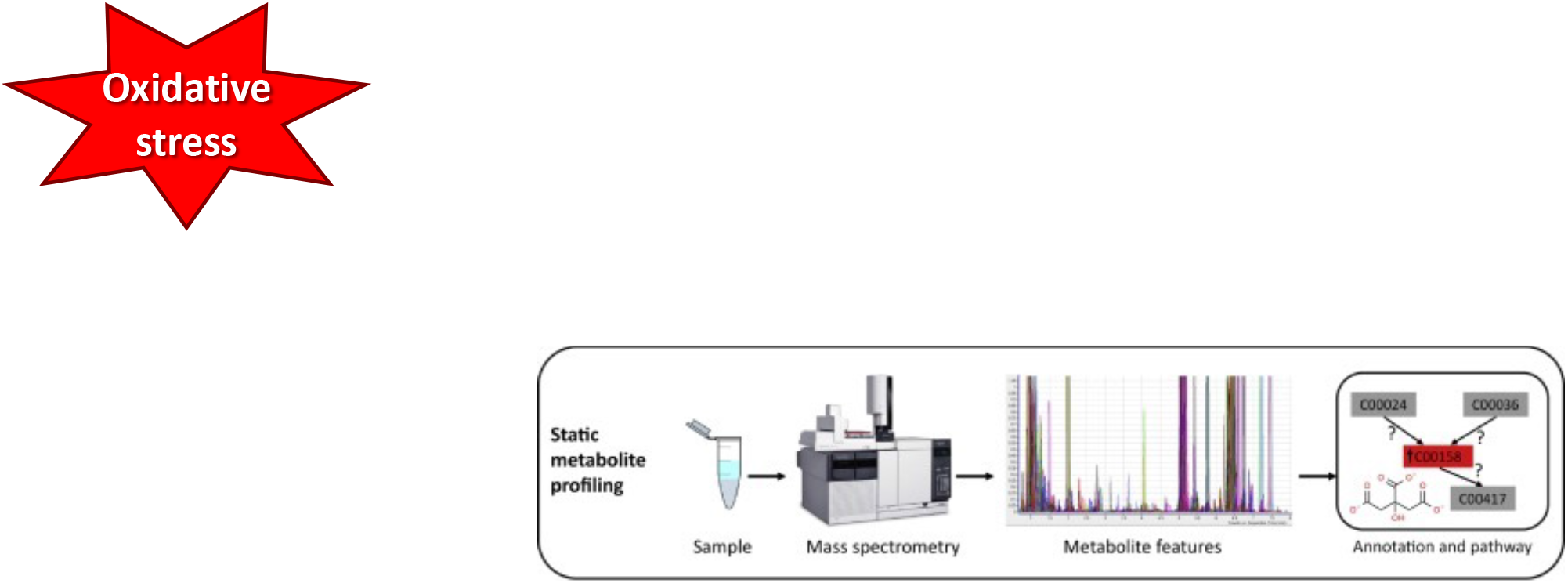

## Introduction

Hypertrophic cardiomyopathy (HCM), the most common cardiac genetic disease worldwide [1], is characterized by myocyte hypertrophy, fibrosis, left ventricular hypercontractility [2], diastolic dysfunction [3], and diverse clinical outcomes, ranging from asymptomatic to heart failure and sudden death [4]. HCM stems from pathogenic variants in sarcomeric protein genes that induce changes in myofilament calcium sensitivity [5–7], crossbridge kinetics [8–10], cardiac mechanics [3, 11] signaling pathways [12] and gene expression [13, 14]. The incorporation of mutant proteins in sarcomeres increases the energy demand for tension development [15] and promotes stress [15], which has led to the notion that HCM is a metabolic disease [16–18].

Metabolic adaptation and structural remodeling in the heart are functionally linked [18–20]. Cardiac hypertrophy is associated with metabolic alterations that promote changes in gene expression. Furthermore, intermediary metabolites in the heart serve as signals to initiate and sustain cardiac adaptation to stress [21]. Hence, understanding metabolic alterations induced by pathogenic variants of sarcomeric proteins may advance the design of metabolism-based therapies [22], preventing the development and progression of HCM.

Recent advances in untargeted mass spectrometry-based metabolomics have expanded our understanding of the mechanistic links between metabolism and the development and progression of human diseases [23–26]. Studies in HCM patients at established disease stages have demonstrated *genotype-independent* mitochondrial dysfunction and broad metabolic/lipid remodeling in heart tissue [27–29]. However transcriptomic and functional studies in 2 mouse models (R92W-TnT, R403Q-MyHC) at the early disease stage revealed *genotype-specific* changes in the expression of metabolic genes, mitochondrial function, and cellular redox state [13]. It is currently unknown whether a common cardiac metabolic trait is present in HCM hearts, at the early-disease stage. To address this question and define the underlying biochemical pathology in early-stage HCM, we conducted cardiac phenotyping by echocardiography, in-depth untargeted mass spectrometry (MS)-based metabolomics and lipidomics analysis followed by pathway and network analysis, and mathematical modeling [30] in R92W-TnT [9, 31] and R403Q-MyHC [32] mouse hearts [13, 14]. We observed allele-specific cardiac metabolic and lipidomic remodeling and differences in substrate utilization and energy provision, which could underlie differences in cardiac HCM phenotype at early disease stage.

## Results

### Pathogenic variants in *TnnT2* and *Myh6* promote gene expression changes at early disease stage

We used two well characterized murine models of HCM [8, 9, 13, 14, 31, 33–35] that express pathogenic variants in two different sarcomeric proteins, Troponin T (TnT*; Tnnt2* gene) or α-Myosin Heavy Chain (MyHC; *Myh6* gene) (**Figure 1A** and **1B**) to define the metabolic phenotype in HCM hearts, at early disease stage (5 weeks of age). Data from mutant hearts was compared to respective littermate control hearts, which we refer to as wild type (WT). Using echocardiography, we observed significantly lower left ventricle (LV) mass and smaller LV cavity size in mutant TnT hearts, and a trend towards higher LV mass in mutant MyHC hearts, when compared to respective WT (**Figure 1C**). Only TnT mutants had evidence of diastolic dysfunction, manifested as lower mitral early [E] and late [A] velocities. (**Figure 1C** and **D**). To dissect genomic heterogeneity and explore the regulatory changes induced by pathogenic variants in TnT and MyHC, we examined our previously published RNA-seq data from 5-week-old mouse hearts [13].

**Figure 1.**
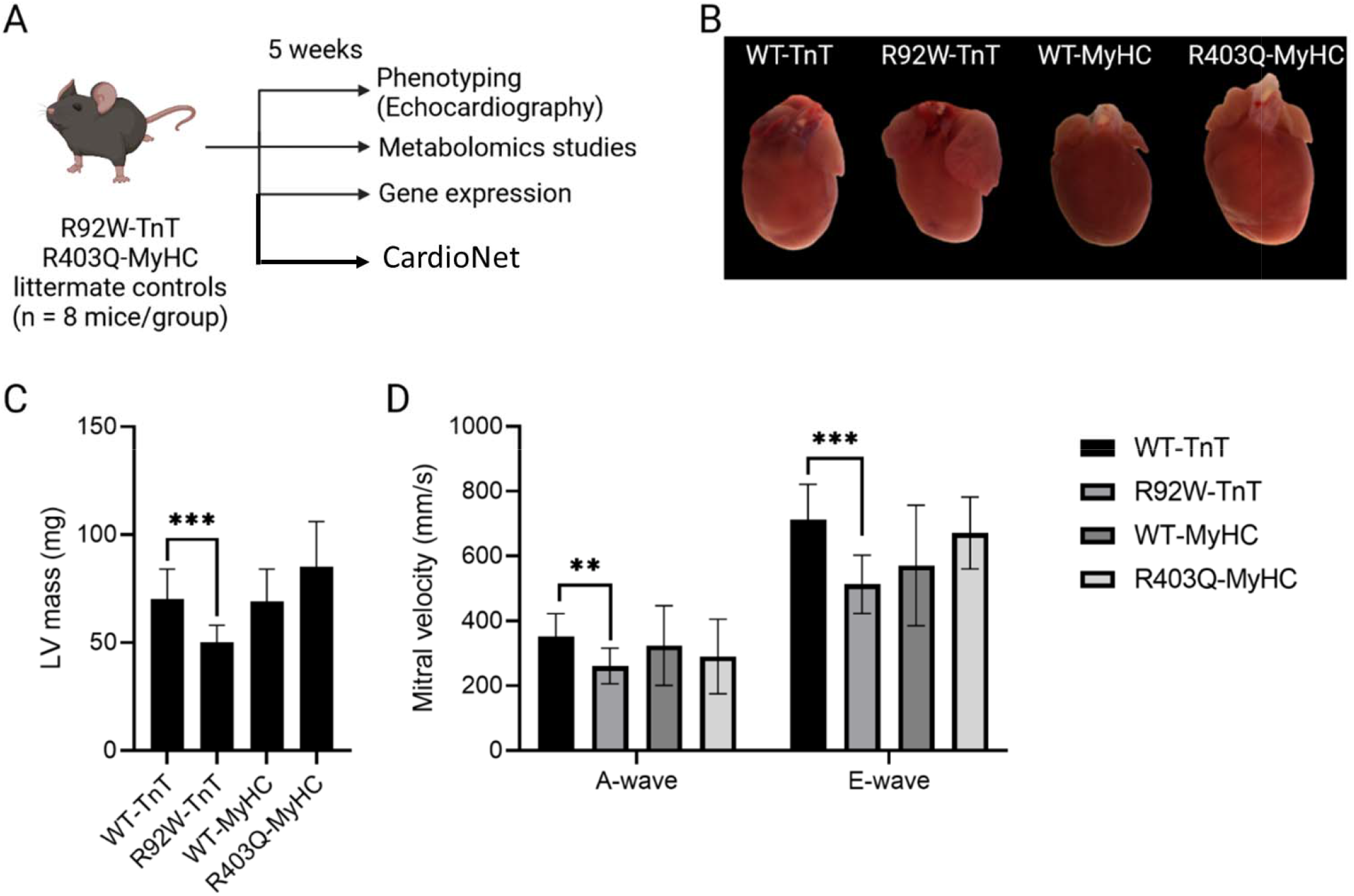
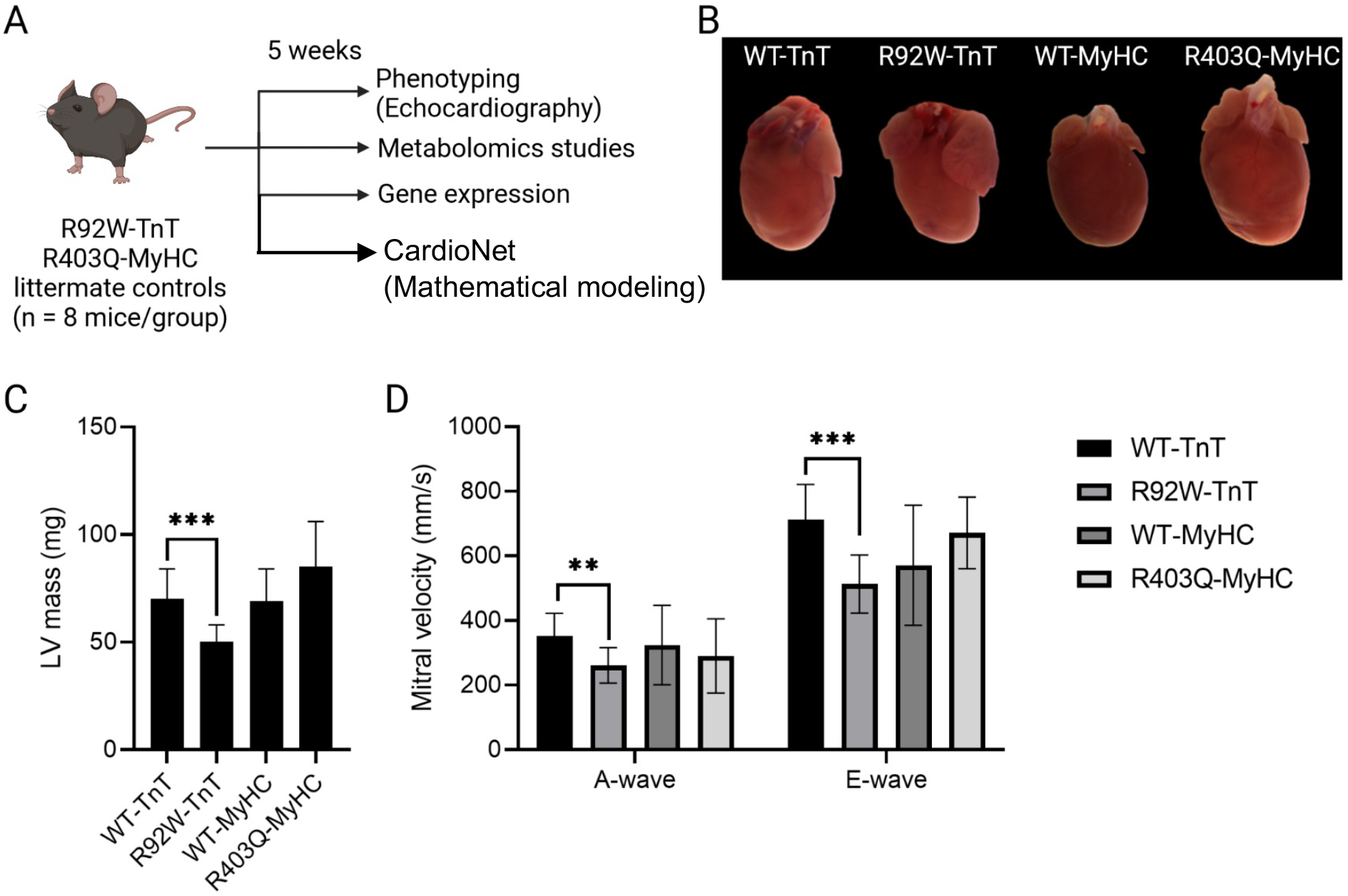
Cardiac remodeling in 2 murine models of HCM at early disease stage. (**A**) Study design: Phenotyping and multi-omics in R92W-TnT and R403Q-MyHC mutant hearts and littermate controls at 5 weeks of age. Hearts were collected for the extraction of primary metabolites, complex lipids, and biogenic amines. RNA-seq analysis, and untargeted metabolomics analysis using three different mass spectrometry (MS) platforms was followed by multi-omics network analysis and computer simulations, using CardioNet. (**B**) Representative gross anatomy from (R92W-TnT, R403Q-MyHC) mice and littermate controls (WT). Bi-arterial enlargement in R92W-TnT mutant heart is visible. (**C**-**D**) Echocardiographic assessment of R92W-TnT and R403Q-MyHC mutant mice. Left ventricular (LV) mass (**C**) and mitral velocity (**D**) indicates early remodeling. n = 8 mice/ group. Wilcoxon rank sum test and rank sum exact test. *P*-value **<0.01; *P*-value ***<0.001.

Principal component analysis (PCA) revealed distinct gene expression patterns in mutant hearts (**Supplemental Figure S1**). Compared to WT, a total of 9,493 and 7,408 genes were upregulated in TnT and MyHC mutants, respectively, with 4,780 genes overlapping between the two mutants (p < 0.05, q < 0.01, FDR of 1%). In contrast, 6,288 and 8,122 genes were down-regulated in TnT and MyHC mutants, respectively. For functional annotation and integrative analysis, we utilized both Reactome [36] and Gene Expression Omnibus (GEO) [37, 38] databases to comprehensively assess differentially expressed protein-coding genes (DEGs) in R92W-TnT and R403Q-MyHC mutant hearts. We constructed enrichment plots for biological processes (**Figures 2A** and **2B**), and interactions between biological processes involved in DEGs were clustered according to their gene and pathway interactions (**Figures 2C** and **2D**). Functional enrichment analysis showed that many DEGs in TnT mutants were predominantly enriched in the Krebs cycle, respiratory electron transport, and branched-chain amino acid metabolism (**Figure 2A)**. In contrast, MyHC mutants primarily showed enrichment in liver-X-receptor (LXR) signaling, mitochondrial biogenesis, and calcium homeostasis (**Figure 2B**). Taken together, these distinct transcription profiles suggest allele-specific metabolic changes.

**Figure 2.**
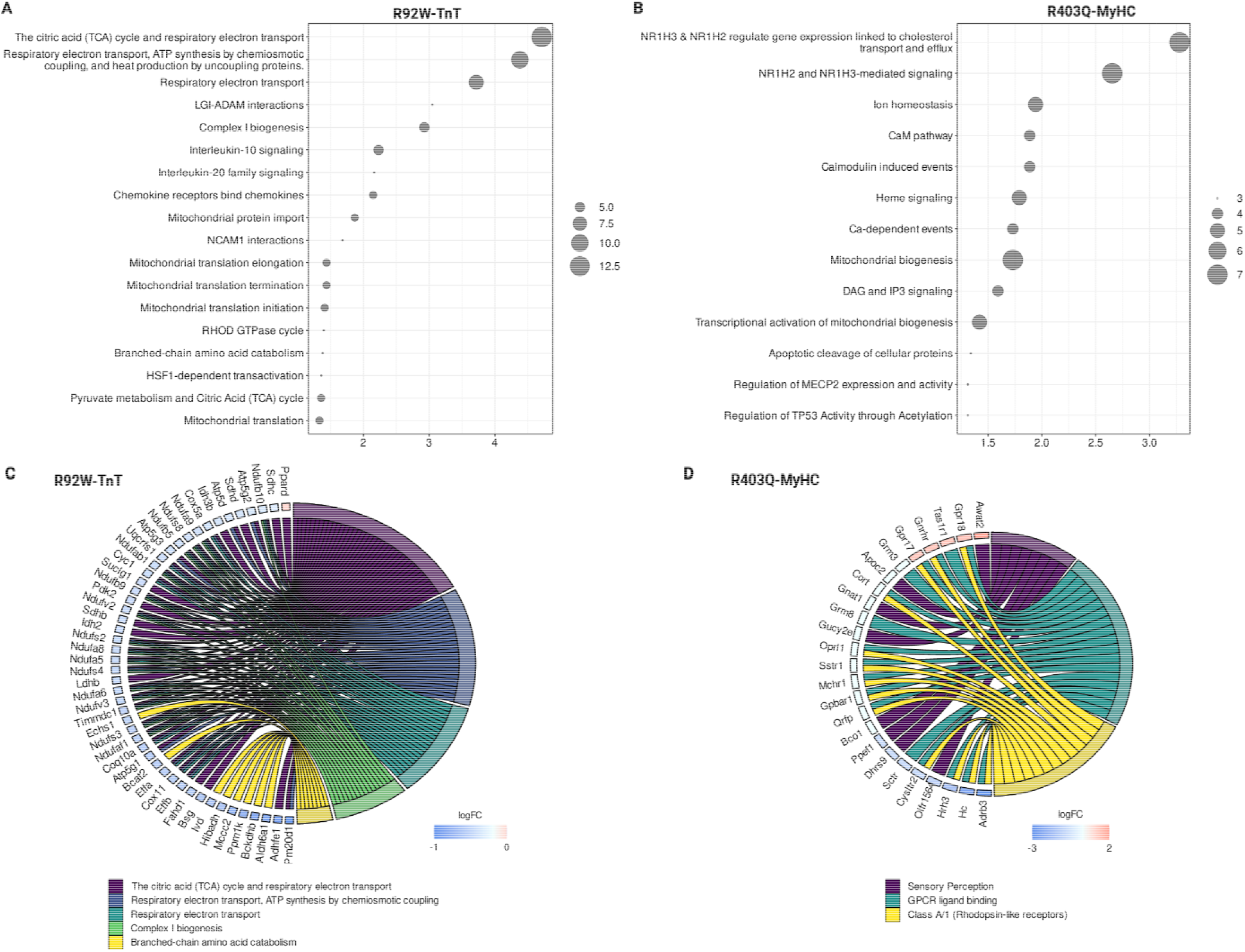
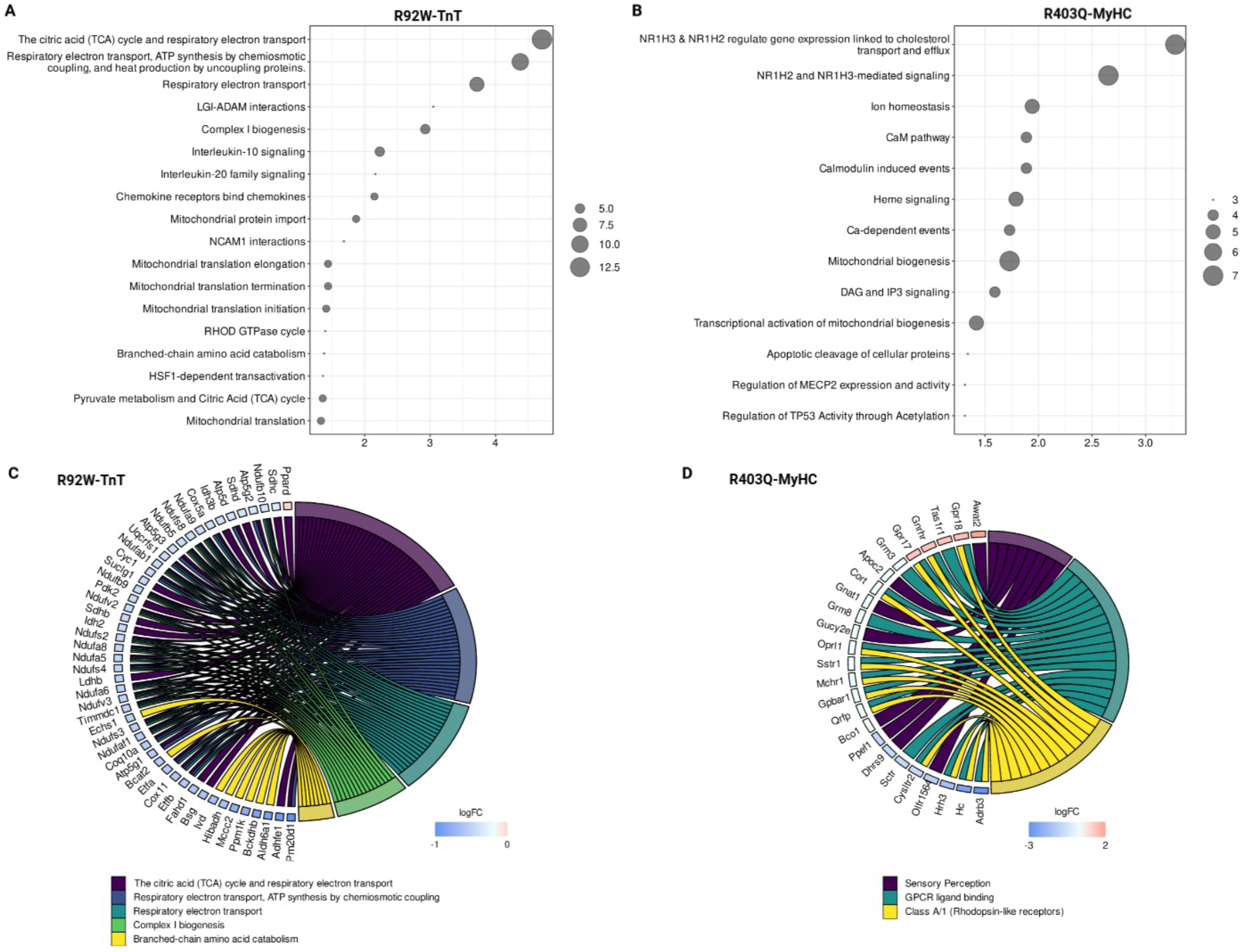
Functional enrichment analysis of differentially expressed protein-coding genes in R92W-TnT and R403Q-MyHC mouse hearts, at early disease stage. (**A**-**B**) GEO database enrichment dot of significantly up- or down-regulated genes in response to (**A**) R92W-TnT or (**B**) R430Q-MyHC mutation. Enrichment for GEO biological processes are depicted against p-value of expression. The dot size represents the number genes under a specific term. (**C**-**D**) GO cluster plot showing a chord dendrogram of the clustering of the expression spectrum of significantly up-or-downregulated genes in response to (**C**) R92W-TnT or (**D**) R430Q-MyHC variants.

### Pathogenic variants in *TnnT2* and *Myh6* differentially alter cardiac lipid composition at early disease stage

To quantify metabolic alterations that could drive cardiac remodeling, we conducted an in-depth untargeted mass spectrometry (MS)-based metabolomics and lipidomics analysis (**Figure 3A**) followed by pathway and network analysis to determine biological patterns [23–25, 39]. Primary metabolites, biogenic amines, and lipids differ in their chemical properties; thus, we combined different chromatographic approaches to increase coverage of metabolomic detection. We identified 891 metabolites in total (**Figure 3A**), including 163 primary metabolites, 165 biogenic amines, and 558 complex lipids using a combination of gas chromatography (GC)-time of flight (TOF) MS, hydrophilic interaction chromatography (HILIC) TripleTOF MS/MS and charged surface hybrid (CSH) chromatography coupled to a quadrupole TOF MS/MS (CSH-QTOF MS/MS), respectively (**Supplemental Table 1-3**). The variability between HCM models was analyzed by principal component analysis (PCA). Among mutant-dependent separation, 64% of the total variance was captured in the first two dimensions across R92W-TnT and R403Q-MyHC mice (**Figure 3B**). The CSH-QTOF MS/MS analysis yielded a total of 514 unique lipid identifications for the MyHC and TnT cohorts across five lipid categories: phospholipids, sphingolipids, glycerolipids, fatty acids (FA) and sterols (**Figure 3C**). Identified lipids comprised 237 phospholipids, 170 glycerolipids, 60 sphingolipids, 4 sterols and 43 fatty acids across the TnT and MyHC cohort. The breakdown of lipid category designation into classes showed 5 phospholipids (phosphatidylinositols (PI), phosphatidylcholines (PC), phosphatidylethanolamines (PE), phosphatiylserine (PS), and cardiolipins (CL)), 3 sphingolipids (sphingomyelins (SM), ceramides (Cer) and N-acylsphingosines (Cer-NS)), 3 glycerolipids (triacylglycerolipids (TG), diacylglycerolipids (DG), and phosphatidylglycerol (PG)), 2 sterols (cholesterol and cholesteryl ester (CE)) and fatty acids (free fatty acids and acylcarnitines) (**Figure 3C**).

**Figure 3.**
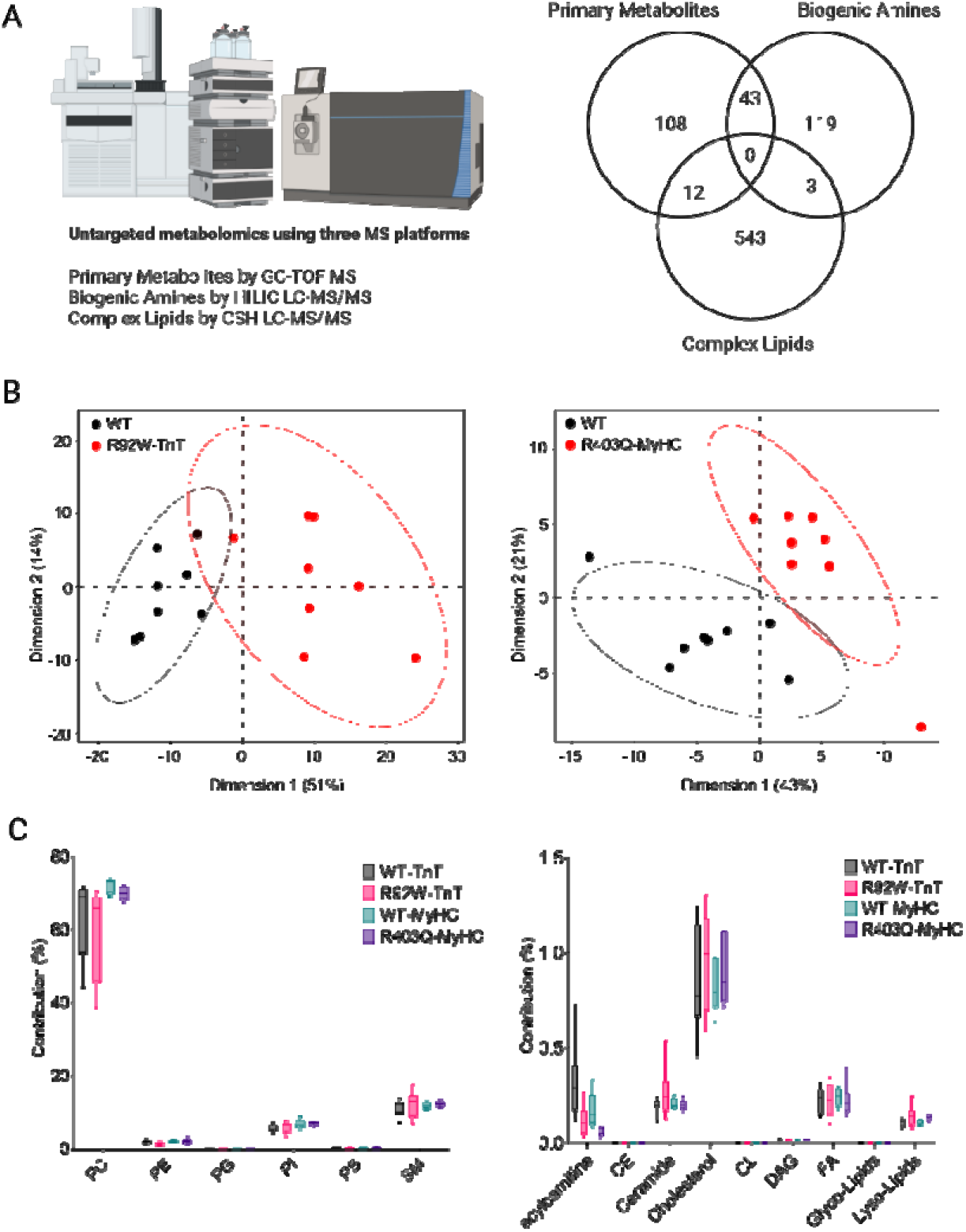
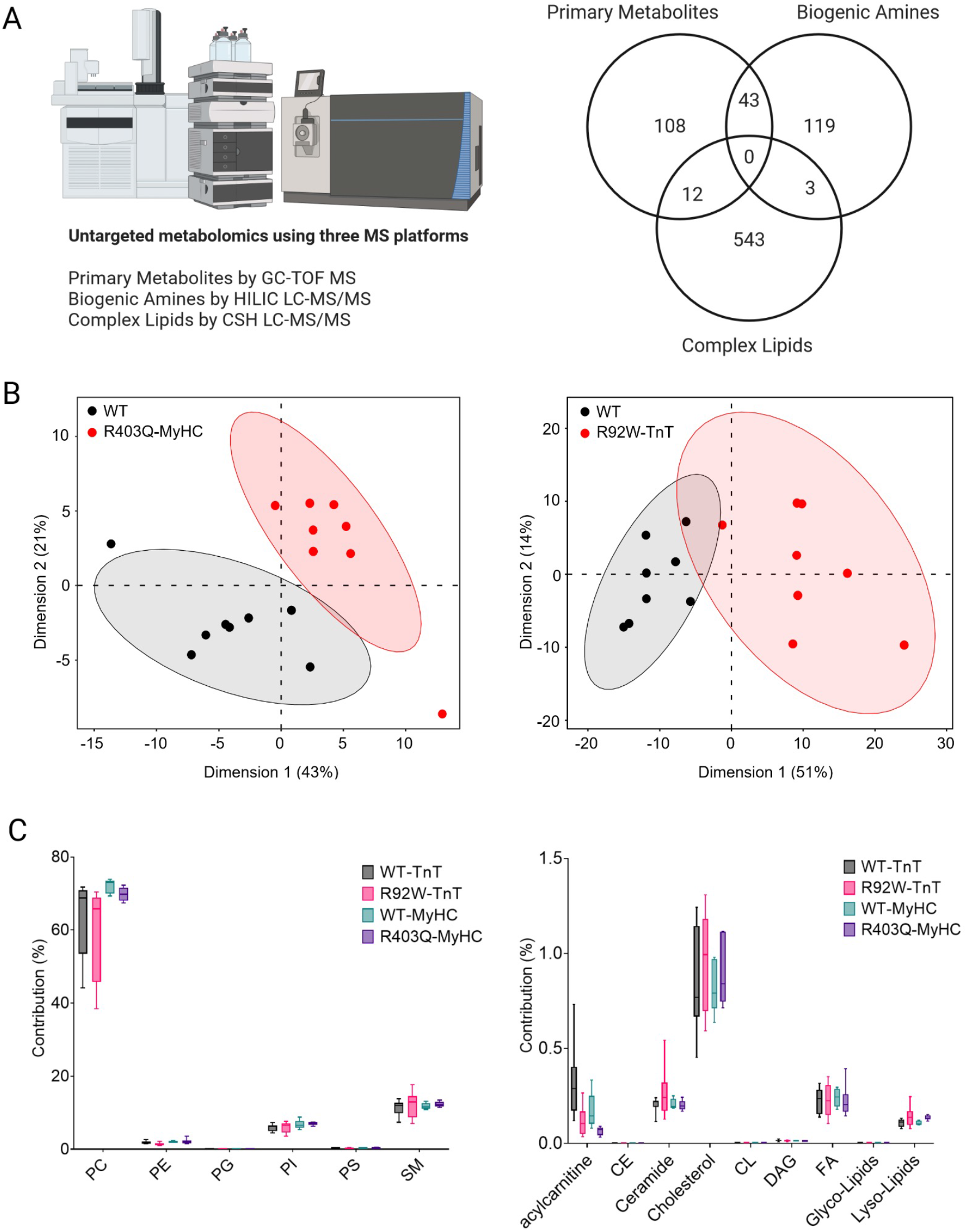
Metabolic and lipidomic profiling R92W-TnT and R403Q-MyHC mutant mouse hearts, at early disease stage. (**A**) Three different mass spectrometry (MS) platforms were used for untargeted metabolomics and lipidomics of heart tissue in R92W-TnT and R403Q-MyHC mouse models. A total of 891 metabolites; with 163 total metabolites found in gas chromatography (GC)-time of flight (TOF) MS, 165 metabolites found in liquid chromatography (LC)-MS/MS analysis, and 563 metabolites found in charged surface hybrid (CSH) LC-MS/MS lipidomics. (**B**) Principal component analysis of the metabolome and lipidome from heart tissue in R92W-TnT and R403Q-MyHC mice. Dimensions 1 and 2 explain 64% of the variance in both genotypes. (**C**) Phospholipid composition in MyHC and TnT mutants. Contribution of phospholipid species is depicted in relation to the total phospholipid fraction. Abbreviations: SM, sphingomyelin; PC, phosphatidylcholine; PE, Phosphatidylethanolamine; PG, phosphatidylglycerol; PI, phosphatidylinositol.

We compared cardiac metabolites of R92W-TnT and R403Q-MyHC mice with WT, using Chemical Similarity Enrichment Analysis (ChemRICH) to identify differences in lipid and intermediary metabolite profiles.^40^ Our analysis identified sets of metabolites based on their chemical ontologies and structural similarity (**Figure 4A** and **B**), and results were visualized according to the overall chemical diversity and enrichment across all metabolites. We observed a stronger biological response in TnT-mutant hearts when compared to MyHC mutants: 40.4% (271/670) of lipids and 24.7% (81/328) of metabolites were significantly different (p<0.05) in TnT mutants compared to WT, whereas 10.4% (70/670) of lipids and 5.8% (19/328) of metabolites were significantly different (p<0.05) in MyHC mutants compared to WT. Both mutant hearts demonstrated significant alterations in purine nucleoside, trisaccharide levels and dicarboxylic acids, as well as lipid species including, acylcarnitines, phosphatidylethanolamines (PEs), phosphatidylinositols (PIs), ceramides (Cer), and triglycerides (TGs) (**Figure 4A** and **4B**). The R92W-TnT variant was associated with reduced abundance of nicotinamide (pyridines), adenine (purines), guanosine (purine nucleosides), leucine (branched-chain amino acids), alpha ketoglutarate (Krebs cycle intermediate) indicating impaired energy provision, compared to littermate controls (**Table 1, Supplemental Figure S2A**). In addition, R92W-TnT hearts demonstrated significantly altered 12 lipid metabolite sets with unsaturated PCs, unsaturated PEs, and unsaturated Cer on the one hand, and lower significance for saturated PCs, saturated lyso-PCs, and saturated Cer on the other hand (**Supplemental Figure S2A**). Additional diversity within phospholipids was observed in the fatty acid linkages, including alkenyl ether (plasmalogen, P-), alkyl ether (O-) and lysoacyl species (e.g., N-acyl-lyso and lyso species), as well as the glycosylation of sphingolipids. Of the identified lipids, marked difference was observed in statistical significance for acylcarnitine, ceramide (Cer) and cardiolipin (CL) levels (**Supplemental Figure S2A,** **Table 1**A). Our data demonstrate cardiac metabolic and lipidomic remodeling in response to pathogenic variants in *TnnT2* and *Myh6* genes. Furthermore, our data suggests that the R92W-TnT variant leads to significant impairment of oxidative metabolism that causes broad phospholipid alterations.

**Table 1A.**
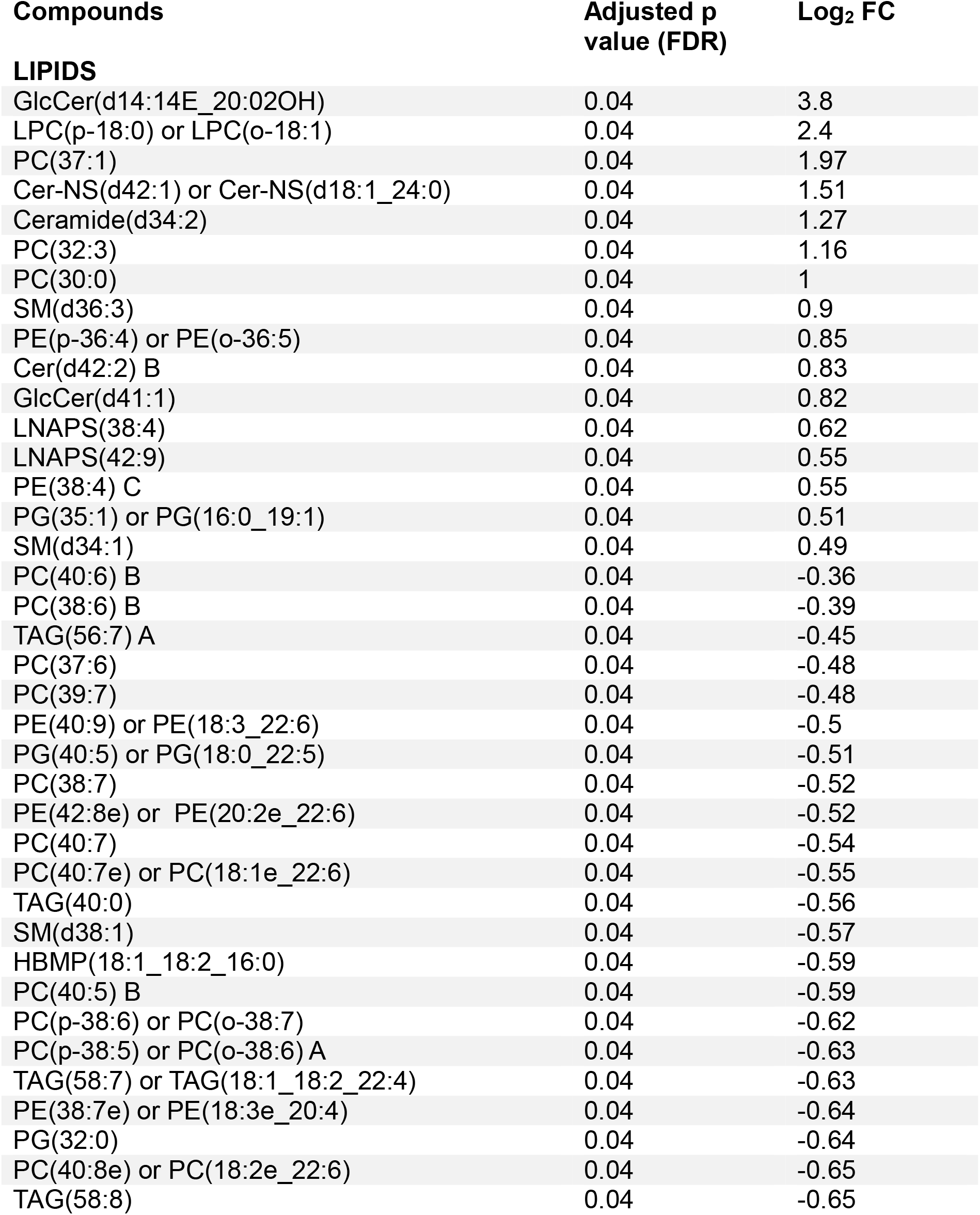
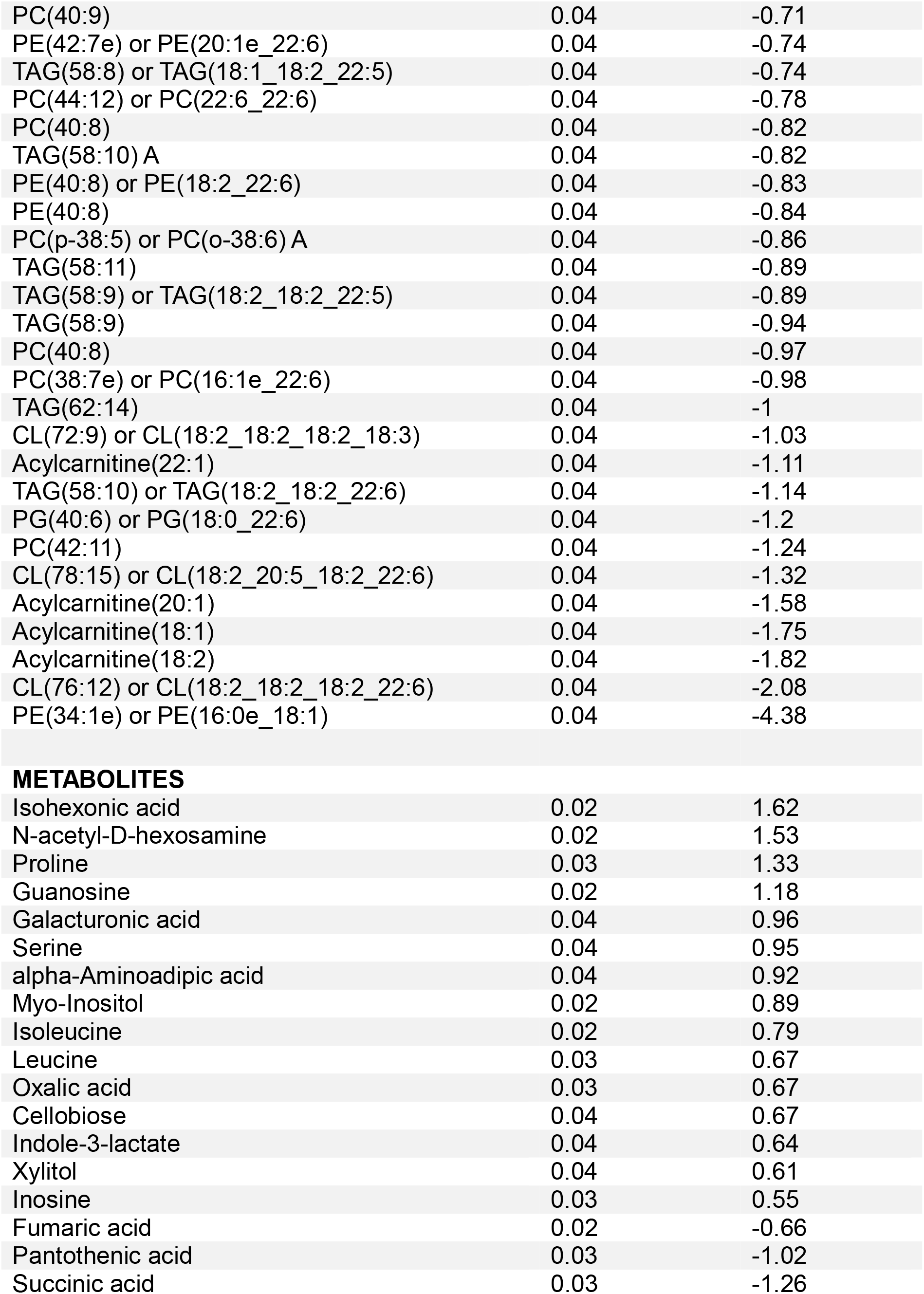
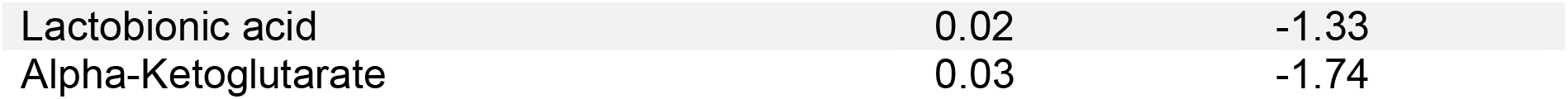
R92W-TnT mutant vs littermate control (WT) mice: differentially expressed Metabolites and Lipids.

| Compounds | Adjusted p value (FDR) | Log<sub>2</sub> FC |
| --- | --- | --- |
| **LIPIDS** |  |  |
| GlcCer(d14:14E_20:02OH) | 0.04 | 3.8 |
| LPC(p-18:0) or LPC(o-18:1) | 0.04 | 2.4 |
| PC(37:1) | 0.04 | 1.97 |
| Cer-NS(d42:1) or Cer-NS(d18:1_24:0) | 0.04 | 1.51 |
| Ceramide(d34:2) | 0.04 | 1.27 |
| PC(32:3) | 0.04 | 1.16 |
| PC(30:0) | 0.04 | 1 |
| SM(d36:3) | 0.04 | 0.9 |
| PE(p-36:4) or PE(o-36:5) | 0.04 | 0.85 |
| Cer(d42:2) B | 0.04 | 0.83 |
| GlcCer(d41:1) | 0.04 | 0.82 |
| LNAPS(38:4) | 0.04 | 0.62 |
| LNAPS(42:9) | 0.04 | 0.55 |
| PE(38:4) C | 0.04 | 0.55 |
| PG(35:1) or PG(16:0_19:1) | 0.04 | 0.51 |
| SM(d34:1) | 0.04 | 0.49 |
| PC(40:6) B | 0.04 | -0.36 |
| PC(38:6) B | 0.04 | -0.39 |
| TAG(56:7) A | 0.04 | -0.45 |
| PC(37:6) | 0.04 | -0.48 |
| PC(39:7) | 0.04 | -0.48 |
| PE(40:9) or PE(18:3_22:6) | 0.04 | -0.5 |
| PG(40:5) or PG(18:0_22:5) | 0.04 | -0.51 |
| PC(38:7) | 0.04 | -0.52 |
| PE(42:8e) or PE(20:2e_22:6) | 0.04 | -0.52 |
| PC(40:7) | 0.04 | -0.54 |
| PC(40:7e) or PC(18:1e_22:6) | 0.04 | -0.55 |
| TAG(40:0) | 0.04 | -0.56 |
| SM(d38:1) | 0.04 | -0.57 |
| HBMP(18:1_18:2_16:0) | 0.04 | -0.59 |
| PC(40:5) B | 0.04 | -0.59 |
| PC(p-38:6) or PC(o-38:7) | 0.04 | -0.62 |
| PC(p-38:5) or PC(o-38:6) A | 0.04 | -0.63 |
| TAG(58:7) or TAG(18:1_18:2_22:4) | 0.04 | -0.63 |
| PE(38:7e) or PE(18:3e_20:4) | 0.04 | -0.64 |
| PG(32:0) | 0.04 | -0.64 |
| PC(40:8e) or PC(18:2e_22:6) | 0.04 | -0.65 |
| TAG(58:8) | 0.04 | -0.65 |
| PC(40:9) | 0.04 | -0.71 |
| PE(42:7e) or PE(20:1e_22:6) | 0.04 | -0.74 |
| TAG(58:8) or TAG(18:1_18:2_22:5) | 0.04 | -0.74 |
| PC(44:12) or PC(22:6_22:6) | 0.04 | -0.78 |
| PC(40:8) | 0.04 | -0.82 |
| TAG(58:10) A | 0.04 | -0.82 |
| PE(40:8) or PE(18:2_22:6) | 0.04 | -0.83 |
| PE(40:8) | 0.04 | -0.84 |
| PC(p-38:5) or PC(o-38:6) A | 0.04 | -0.86 |
| TAG(58:11) | 0.04 | -0.89 |
| TAG(58:9) or TAG(18:2_18:2_22:5) | 0.04 | -0.89 |
| TAG(58:9) | 0.04 | -0.94 |
| PC(40:8) | 0.04 | -0.97 |
| PC(38:7e) or PC(16:1e_22:6) | 0.04 | -0.98 |
| TAG(62:14) | 0.04 | -1 |
| CL(72:9) or CL(18:2_18:2_18:2_18:3) | 0.04 | -1.03 |
| Acylcarnitine(22:1) | 0.04 | -1.11 |
| TAG(58:10) or TAG(18:2_18:2_22:6) | 0.04 | -1.14 |
| PG(40:6) or PG(18:0_22:6) | 0.04 | -1.2 |
| PC(42:11) | 0.04 | -1.24 |
| CL(78:15) or CL(18:2_20:5_18:2_22:6) | 0.04 | -1.32 |
| Acylcarnitine(20:1) | 0.04 | -1.58 |
| Acylcarnitine(18:1) | 0.04 | -1.75 |
| Acylcarnitine(18:2) | 0.04 | -1.82 |
| CL(76:12) or CL(18:2_18:2_18:2_22:6) | 0.04 | -2.08 |
| PE(34:1e) or PE(16:0e_18:1) | 0.04 | -4.38 |
| METABOLITES |  |  |
| Isohexonic acid | 0.02 | 1.62 |
| N-acetyl-D-hexosamine | 0.02 | 1.53 |
| Proline | 0.03 | 1.33 |
| Guanosine | 0.02 | 1.18 |
| Galacturonic acid | 0.04 | 0.96 |
| Serine | 0.04 | 0.95 |
| alpha-Aminoadipic acid | 0.04 | 0.92 |
| Myo-Inositol | 0.02 | 0.89 |
| Isoleucine | 0.02 | 0.79 |
| Leucine | 0.03 | 0.67 |
| Oxalic acid | 0.03 | 0.67 |
| Cellobiose | 0.04 | 0.67 |
| Indole-3-lactate | 0.04 | 0.64 |
| Xylitol | 0.04 | 0.61 |
| Inosine | 0.03 | 0.55 |
| Fumaric acid | 0.02 | -0.66 |
| Pantothenic acid | 0.03 | -1.02 |
| Succinic acid | 0.03 | -1.26 |
| Lactobionic acid | 0.02 | -1.33 |
| Alpha-Ketoglutarate | 0.03 | -1.74 |

**Table 1B.** R403Q-MyHC mutant vs littermate control (WT) mice, *p* ≤ 0.01, and *Log_2_ Fold Change* comparisons of Lipids and Metabolites.

| Compounds | FDR $p < 0.05$ | $\text{Log}_2$ FC |
| --- | --- | --- |
| None | - | - |
| **LIPIDS** |  |  |
|  | **Non-adjusted p value** | **$\text{Log}_2$ FC** |
| DAG(38:6) A | 0.005 | -0.31 |
| SM(d40:2) A | 0.010 | -0.43 |
| PC(35:3) | 0.010 | -0.52 |
| DAG(40:8) or DAG(18:2_22:6) | 0.004 | -0.57 |
| PI(34:2) or PI(16:0_18:2) | 0.007 | -0.61 |
| TAG(58:10) A | 0.010 | -0.67 |
| DAG(36:4) A | 0.005 | -0.71 |
| acylcarnitine(10:0) | 0.005 | -0.99 |
| acylcarnitine(14:2) | 0.007 | -1.05 |
| acylcarnitine(20:1) | 0.010 | -1.44 |
| acylcarnitine(18:2) | 0.010 | -1.7 |
| acylcarnitine(18:1) | 0.007 | -1.79 |
| Linoleic acid | 0.010 | -0.65 |
| Oleic acid | 0.005 | -0.83 |
| 3-Hydroxypalmitic acid | 0.004 | -1.21 |
| **METABOLITES** |  |  |
| Maltotriose | 0.001 | 1.04 |
| Guanosine | 0.007 | 0.88 |
| Anserine | 0.010 | -1.66 |

GlcCer is glucosylceramide, LPC is lysophosphatidylcholine, PC is phosphatidylcholine, Cer-NS is ceramide non-hydroxyfatty acid-sphingosine, SM is sphingomyelin, PE is phosphatidylethanolamine, Cer is ceramide, LNAPS is N-acyl-lysophosphatidylserine, PG is phosphatidylglycerol, TAG is triacylglycerol, HBMP is hemibismonoacylglycerophosphate, CL is cardiolipin, DAG is diacylglycerol, and PI is phosphatidylinositol

**Figure 4.**
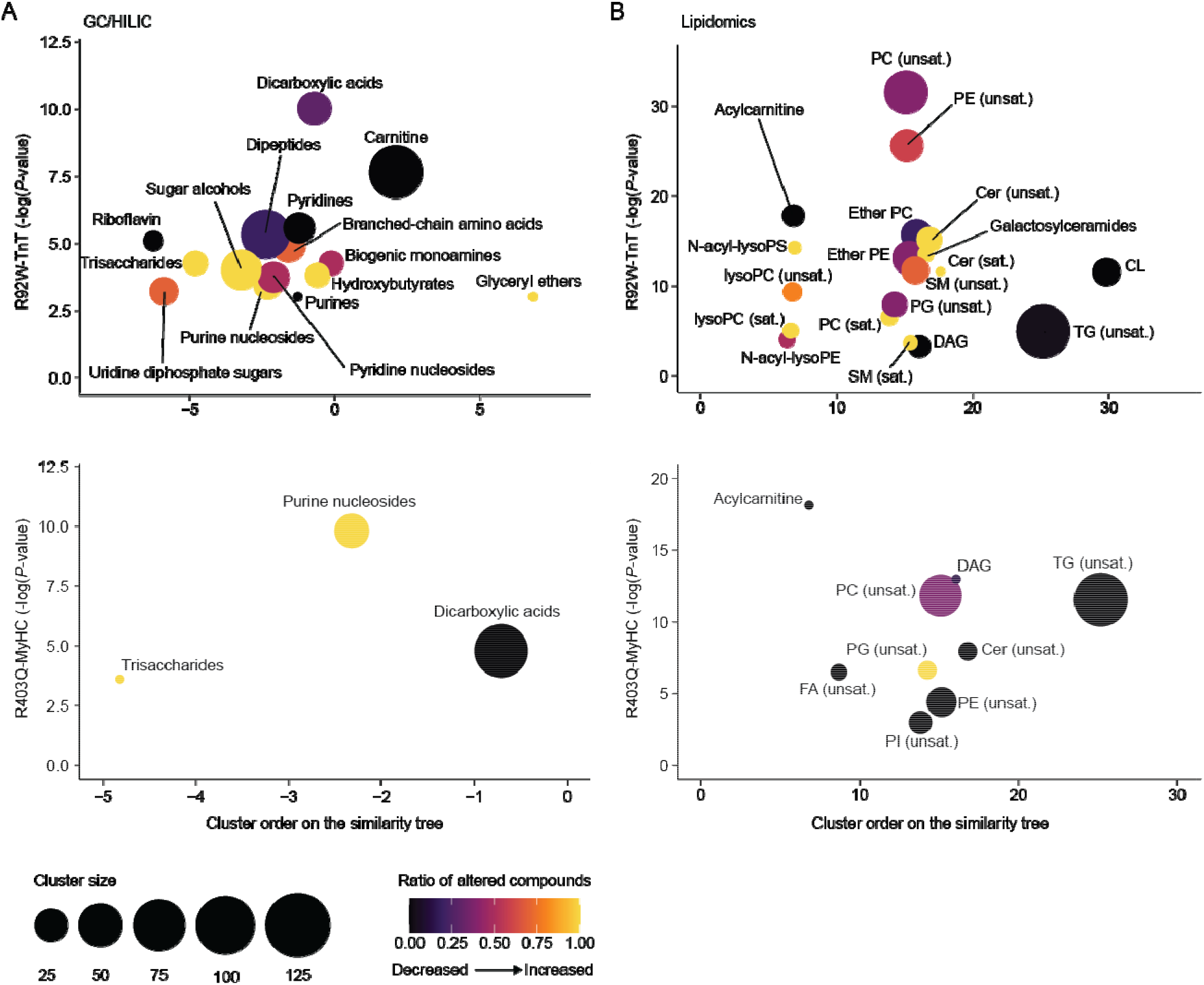
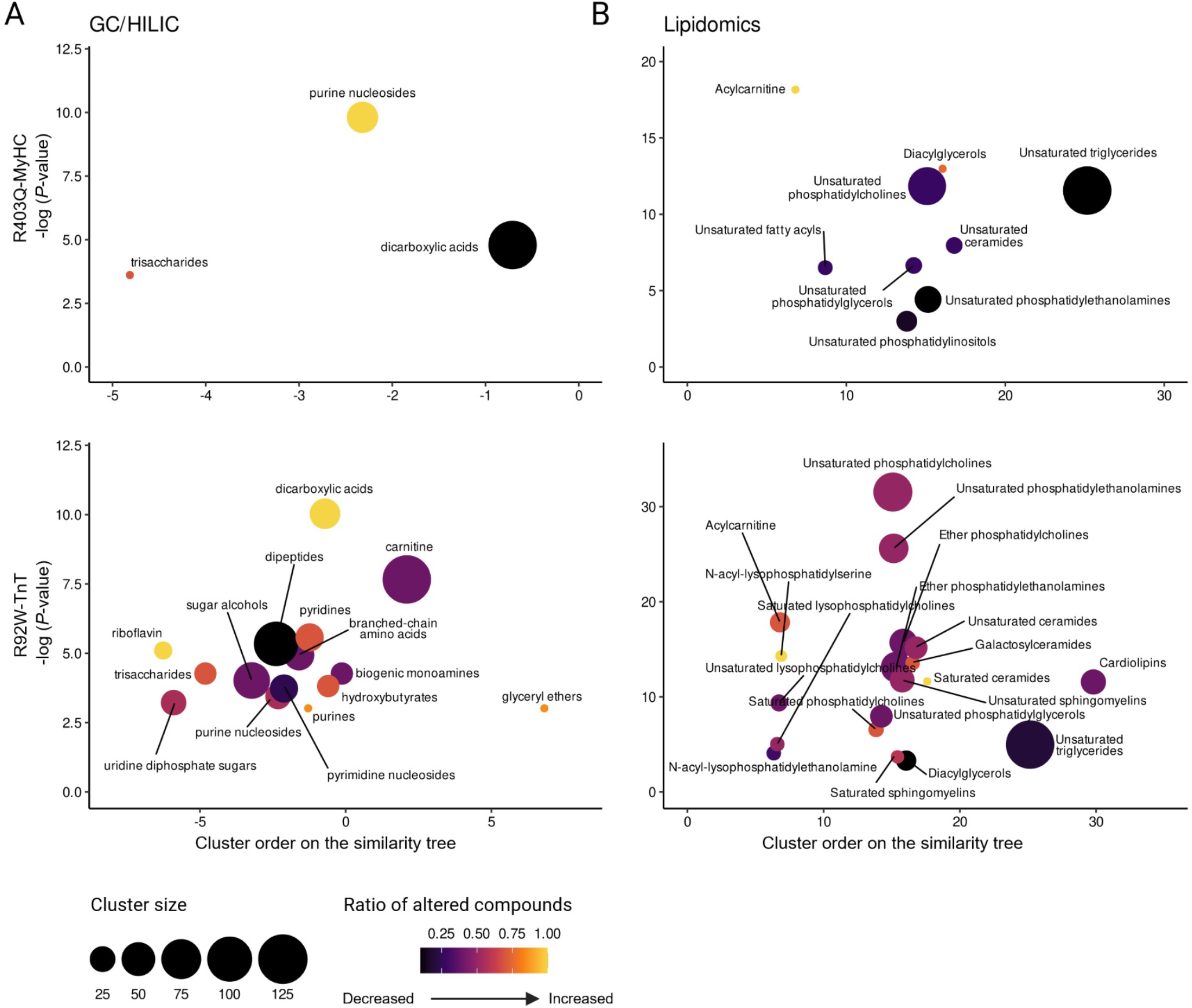
Pathogenic variants in sarcomeric protein genes promote cardiac metabolic remodeling at early disease stage. (**A-B**) ChemRICH set enrichment plots for GC-MS and LC-MS/MS-based metabolomics (**B**) and CSH LC-MS/MS lipidomics (**B**). Each node depicts a significantly altered metabolite cluster. Node sizes and colors represent the total number of metabolites and proportion of increased or decreased compounds in each cluster, respectively. Cer, ceramides; CL, cardiolipins; PC, phosphatidylcholes; PE, phosphatidylethanolamines; PS, phosphatidylserines; SM, sphingomyelins; TG, triglycerides; sat., saturated; unsat., unsaturated

### Pathogenic variants in sarcomeric protein genes impact the acyl-composition of phospholipids in HCM mouse hearts at early disease stage

In a next step, we investigated whether the acyl-chain composition and degree of saturation in acylcarnitine, ceramide and cardiolipin species contribute to the observed difference in metabolic profiles between TnT and MyHC mutant hearts. We quantified a shift in the incorporation of long-chain fatty acyl residues in acylcarnitine, ceramide and cardiolipin species (**Figure 5**). Specifically, we identified lower abundance of unsaturated long-chain acyl-carnitines (18:1, 18:2 and 20:1) in both mutant mouse hearts, compared to WT (**Figure 5A** and **5B**). Next, we compared the fatty acyl chain composition and degree of saturation within the ceramide species **(Figure 5C** and **5D**). Cer d36:1 is the predominant ceramide species (>40%) in hearts from WT mice followed by Cer d40:2 (25-30%) (**Figure 5C**). However, this relationship is shifted towards ceramide d40:2 in both mutants (35% in TnT mutants, 45% in MyHC mutants). The R403Q-MyHC variant is associated with a significantly higher contribution of Cer d40:2 compared to any other experimental group (**Figure 5D**). The increase in fatty acyl chain length and degree of saturation is prominent in TnT mutants as indicated by increased contribution of Cer d40:0 and d42:0, which is unchanged in R403Q-MyHC mutants compared to littermate controls. The CL fraction was significantly impacted in R92W-TnT mice with a dominant shift from CL 76:12 towards CL 72:7 (**Figure 5E** and **5F**). In control mice, CL76:12 and CL72:7 represent on average 18% and 8% of the total CL fraction. This relationship contrasts with TnT mutants, with CL76:12 and CL72:7 representing on average 10% and 15%, respectively, whereas the CL composition was similar in MyHC mutants and WT. Taken together, we observed selective enrichment of FA18:0 in Cer and CL species in TnT mutant hearts, which could underlie differences in cardiac metabolites and mitochondrial function observed between TnT and MyHC mutant hearts at early disease stage.

**Figure 5.**
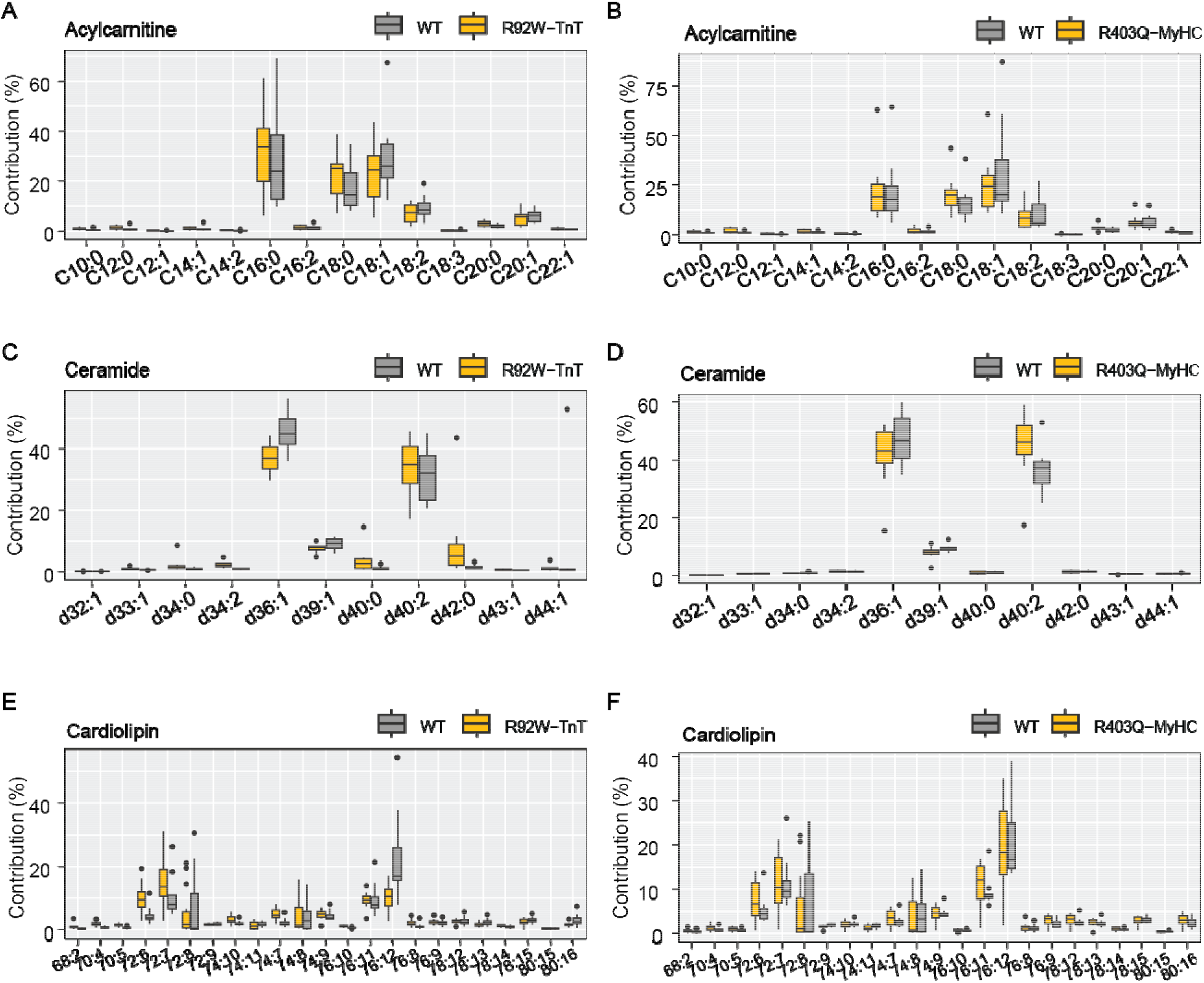
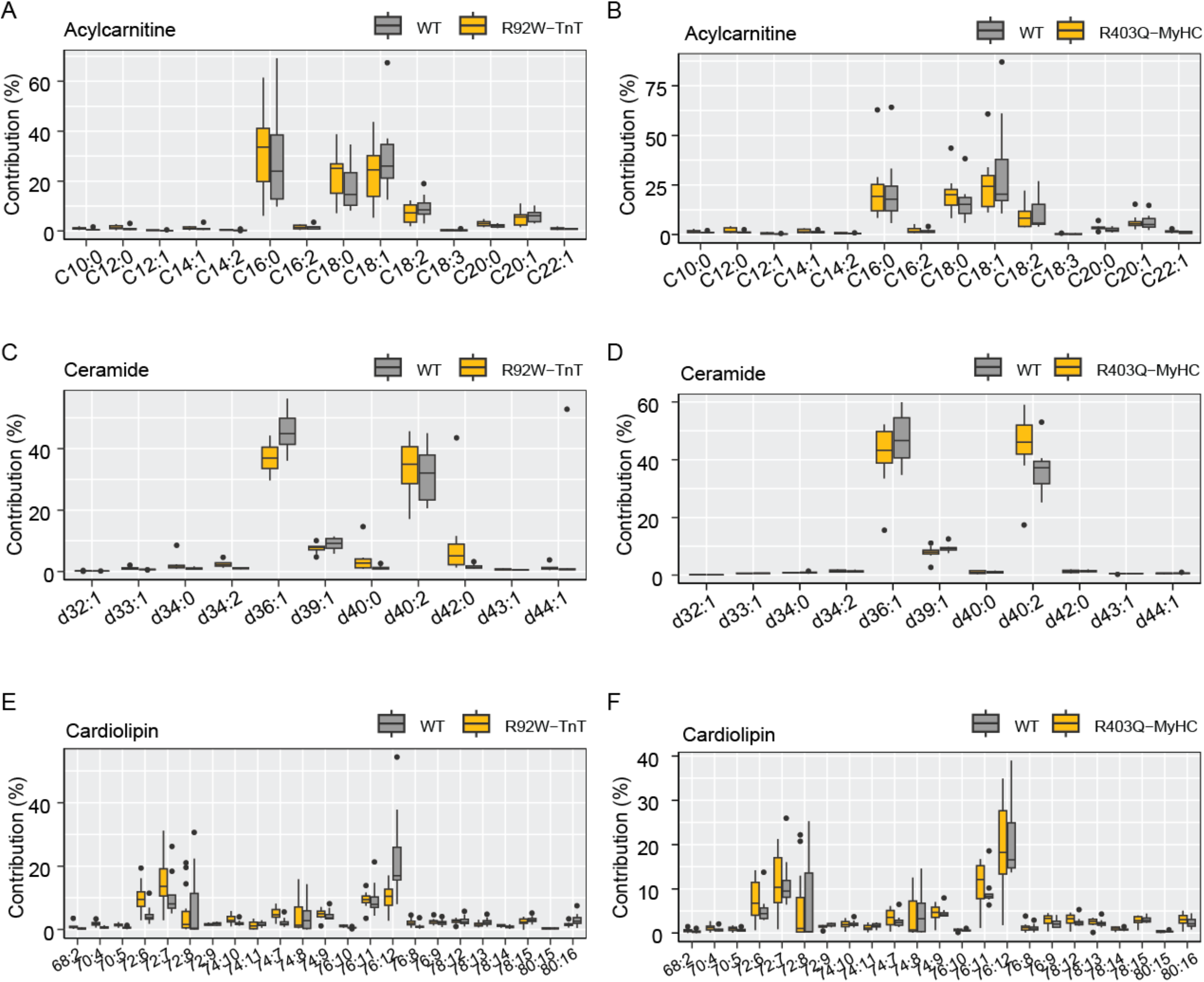
Fatty acyl-chain composition and degree of unsaturation in ceramide, acylcarnitine and cardiolipin species in HCM mouse hearts at early disease stage. (A-B) Acylcarnitine composition in R92W-TnT (A) and R403Q-MyHC (B) hearts compared to littermate controls. (C-D) Ceramide composition in R92W-TnT (C) and R403Q-MyHC (D) hearts compared to littermate controls. (E-F) Cardiolipin composition in R92W-TnT (E) and R403Q-MyHC (F) hearts compared to littermate controls. n = 8-16 sample/group.

### Mathematical flux analysis identifies differential metabolic dependencies in R92W-TnT and R403Q-MyHC mutant hearts at early disease stage

Changes in metabolite and lipid abundances can either result from differential production or utilization of a given intermediate due to higher flux from synthesizing reactions or decreased flux towards consuming reactions. To assess the impact of pathogenic variants in TnT and MyHC on cardiac metabolism, we conducted mathematical modelling using CardioNet [30]. This computational approach allows the prediction and identification of metabolic fluxes by integrating experimental data, including from metabolomics and lipidomics analyses [41, 42]. We determined flux distributions using flux balance analysis (FBA), which applies a steady-state assumption while optimizing an objective function (see **Methods** for details). The linear optimization problem was to maximize ATP provision and ensure biomass synthesis (e.g., protein, lipid synthesis) while fulfilling a set of constraints defined by the metabolic profile in mutant and WT hearts. Annotation enrichment of observed clusters and unsupervised hierarchical cluster analysis (**Supplemental Figure S3A** and **S3B**) demonstrated distinct metabolic remodeling in R92W-TnT and R403Q-MyHC hearts, consistent with our multi-omics analysis. Simulations of TnT mutants revealed upregulation of lipid metabolism as reflected in cluster 2 and cluster 3 (**Supplemental Figure S3A** and S**3B**).

To understand the predicted metabolic flux distributions, we first annotated identified reactions to curated pathways within the CardioNet database. Unsupervised hierarchical clustering of calculated flux rates separated TnT and MyHC mutants from WT mice (**Figure 6A**). We identified an increased flux contribution in fructose, ribose, and glucose metabolism in TnT mutants, which combine glycolysis, oxidative and non-oxidative pentose phosphate pathway, and nucleotide synthesis. Furthermore, we found increased flux distributions in fatty acid biosynthesis and lipid metabolism consistent with our lipidomics data (**Figure 6A**). Overall oxidative phosphorylation was lower in TnT and MyHC mutants compared to WT (**Figure 6A**). To understand the metabolic fate of energy-providing substrates, we analyzed the contribution of glucose and fatty acid utilization. Glucose uptake and utilization is predicted to increase (1.5-fold) in TnT mutants and decrease (0.7-fold) in MyHC mutants when compared to WT (**Figure 6B**). Our simulations also predicted higher uptake of linoleate (FA18:2) and linolenate (18:3) in both mutants, when compared to WT (**Figure 6B**). In TnT mutants, FA18:2 oxidation is increased while FA22:5 and FA22:6 oxidation is decreased compared to control and MyHC mutants (**Figure 6C**). The per carbon ATP yield from FA22:5 and FA22:6 is significantly higher compared to FA18:2 and glucose indicating that MyHC mutant and WT mice rely more on ATP provision from fatty acids compared to TnT mutants. These findings are further supported by network analysis and visualization (**Figure 6D**). Increased lipid remodeling and synthesis in TnT mutants requires compensatory flux changes to ensure adequate provision of reducing equivalents in the form of NADH and NADPH. Taken together, our simulations suggest that energy provision and substrate utilization is differentially impacted in R92W-TnT and R403Q-MyHC mutants. TnT mutants are negatively impacted by a mismatch between oxidative and reductive demands as evidenced by a metabolic nutrient shift from increased fatty acid towards glucose utilization.

**Figure 6.**
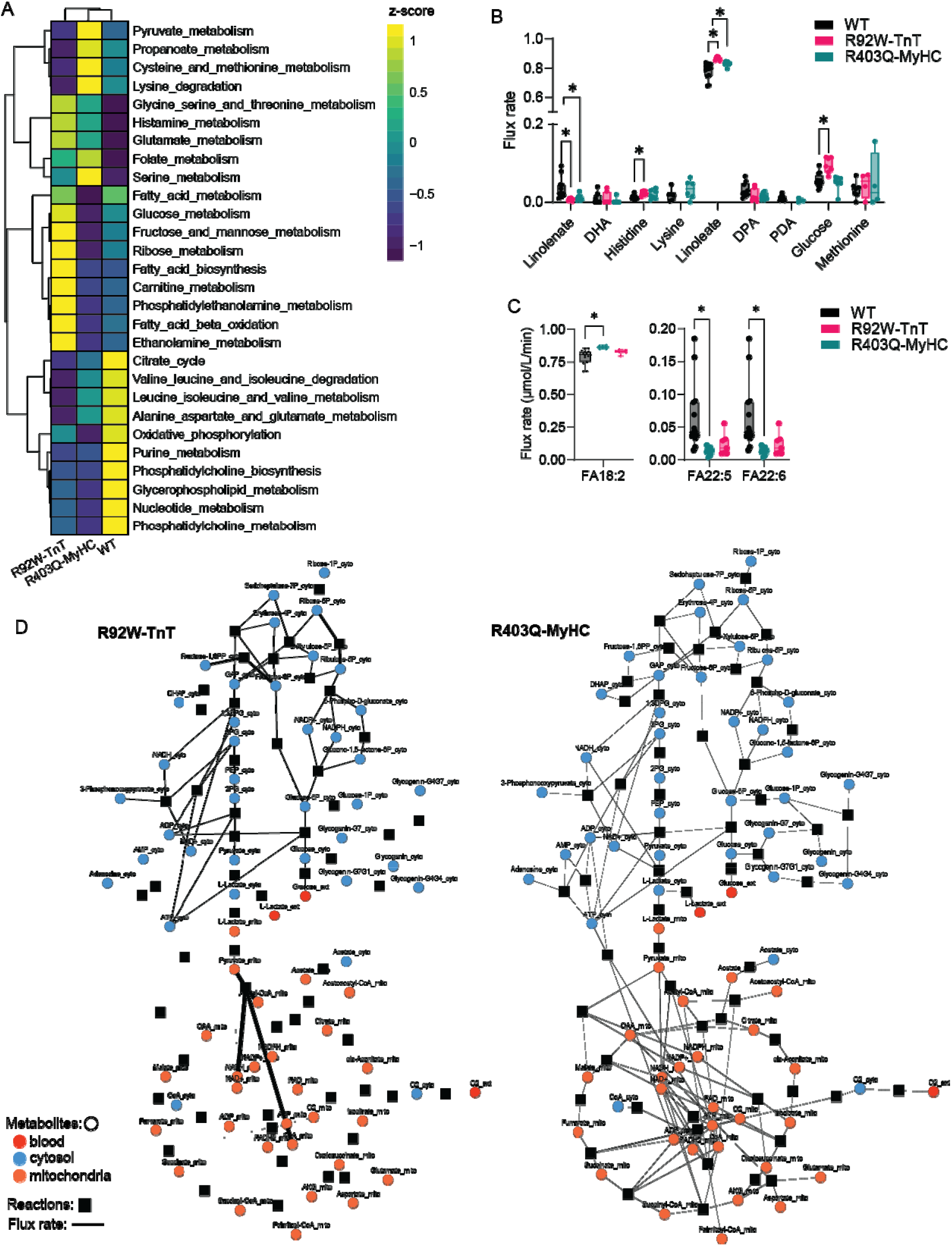
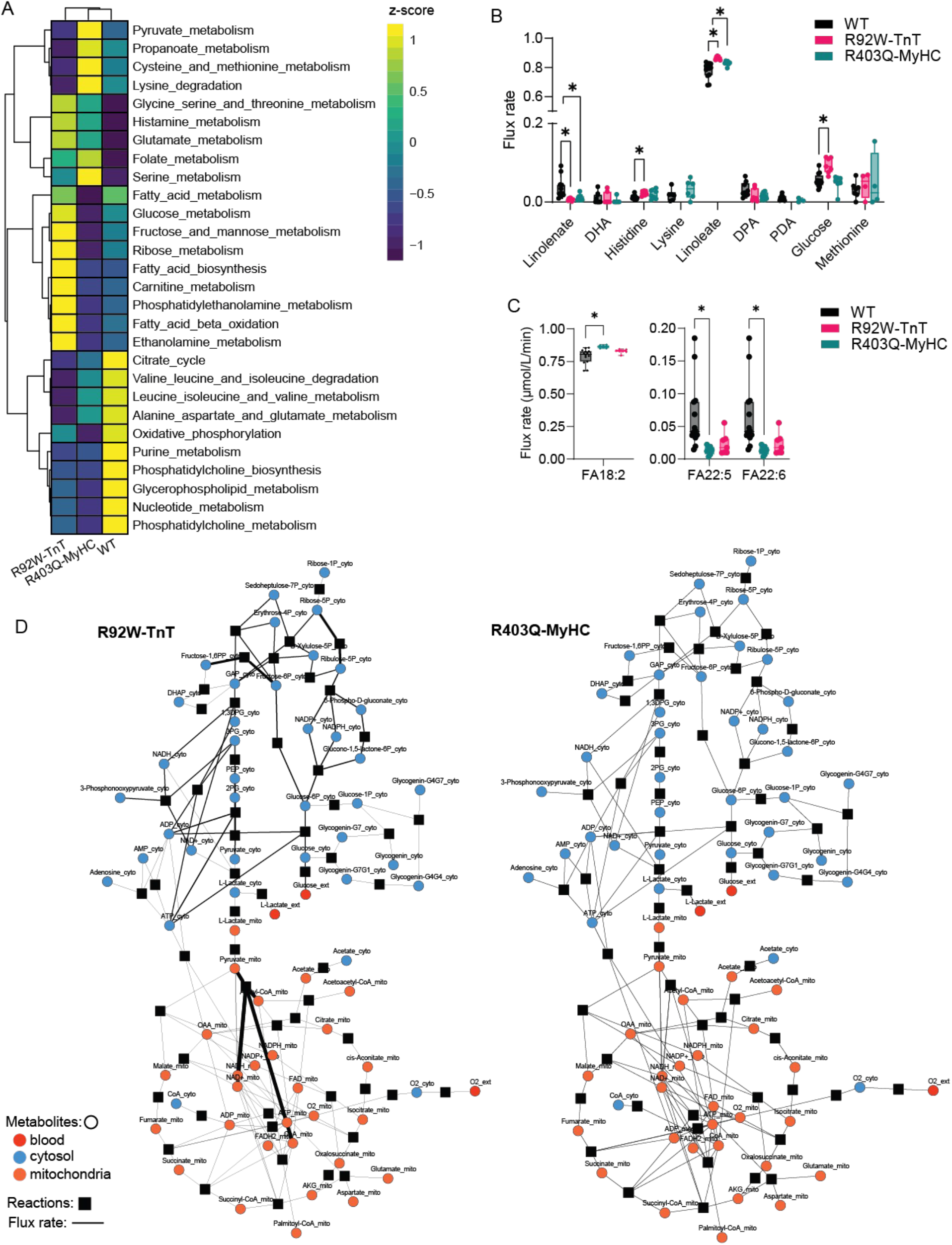
Analysis of flux alterations in R92W-TnT and R403Q-MyHC using CardioNet at early disease stage. (**A**) Metabolic flux distributions were calculated using CardioNet and constrained-base modeling using optimization algorithms. Reactions were assigned to pathways and predicted fluxes were summed across pathways for unsupervised hierarchical cluster analysis. (**B**) Predicted uptake rates of extracellular metabolites across experimental groups. Flux rates were calculated using CardioNet simulations for R92W-TnT, R403Q-MyHC and WT mouse hearts. (**C**) Flux rate predictions for fatty acid β-oxidation for linoleate (FA18:2), docosapentaenoic acid (DPA; FA22:5), and docosahexaenoic acid (DHA, FA22:6) in both mutants compared to WT. (**D**) Visualization of predicted flux rates for glycolysis, pentose phosphate pathway and Krebs cycle. Edge thickness is scaled to calculated flux rates. Simulations n = 14 to 7 per group. ANOVA with multiple comparison analysis. FDR<1%; *q-value<0.05; **q-value<0.01; ***q-value<0.001.

### Allele-specific differences in oxidative stress at early disease stage

Computational modeling revealed increased flux in lipid metabolism. Specifically, CardioNet simulations predicted an increased peroxidation of PS and PE in the lysosome and microsome (combined compartment of endoplasmic reticulum and golgi apparatus) (**Figure 7A**). Increased lipid peroxidation is potentially associated with an increased oxidative stress. To test this hypothesis, we quantified oxidative-induced lipid peroxidation using 4-hydroxynonenal (4-HNE) as a marker protein (**Figure 7B**). 4-HNE is an electrophilic molecule that can easily diffuse across membranes, is highly reactive toward nucleophilic thiol and amino groups, and is a major generator of oxidative stress. TnT mutant hearts had higher abundance of 4-HNE compared to littermate controls, whereas no difference in 4-HNE was observed between MyHC mutants and littermate controls (**Figure 7B**).

**Figure 7.**
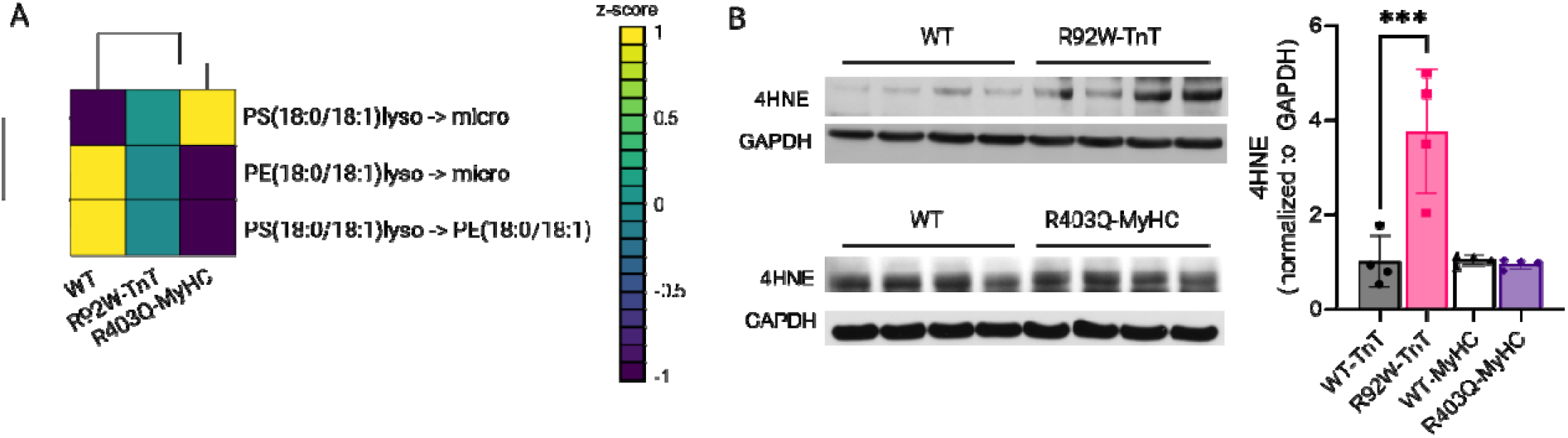
Phospholipid metabolism drives oxidative stress in R92W-TnT hearts. (**A**) Summary of predicted flux rates for modifications of peroxidation of PS and PE in the lysosome (lyso) and microsome (micro; endoplasmic reticulum and golgi apparatus). (**B**) Quantification of lipid peroxidation using 4-hydroxynonenal (4-HNE) and western blotting. Densitometry was normalized to total GAPDH abundance. n=4 mice/group. 2-way ANOVA and Tukey’s posthoc multiple comparisons analysis. FDR<1%; **q-value<0.01

## DISCUSSION

Our analysis of comprehensive untargeted cardiac metabolomics in R92W-TnT and R403Q-MyHC mutant hearts at early disease stage revealed marked differences in metabolic profiles that could provide a mechanistic framework for phenotypic variability in HCM. At early disease stage, R92W-TnT hearts have left atrial (LA) enlargement, diastolic dysfunction, significant impairment of energy substrate metabolism, phospholipid remodeling and evidence of lipid peroxidation, whereas R403Q-MyHC hearts have preserved diastolic function, energy substrate and phospholipid metabolism (**Table 1**, **Figures 1 and 4**). Computer simulations predict upregulation of glycolysis and decreased oxidative phosphorylation in TnT mutants, which could lead to energetic stress, diastolic dysfunction and LA enlargement, features commonly seen in human HCM^43-45^. Interestingly, mutant MyHC hearts showed a trend towards higher LV mass and small changes in the cardiac metabolome and lipidome, suggesting that myocyte hypertrophy and metabolic remodeling occur in parallel in early disease stage of HCM.

We combined RNA-sequencing and metabolomics with computational modeling using CardioNet^41, 42, 46-48^ to identify flux distributions that could explain the observed metabolic patterns. CardioNet predicts an increased contribution of glycolysis and pentose phosphate pathway towards ATP provision and reductive metabolism, along with reduction in oxidative phosphorylation in R92W-TnT hearts (**Figure 6D**). In contrast, mathematical simulations of R403Q-MyHC metabolism indicate that the observed metabolic changes are compensatory flux changes likely protecting R403Q-MyHC hearts (**Figure 6D**). Thus, our systems biology approach using CardioNet reveals allele-specific flux changes, and provide a mechanistic understanding of metabolomics studies.

In humans, the R92W-TnT variant is associated with mild cardiac hypertrophy and sudden death at a young age [49], whereas the R403Q-MyHC variant is associated with moderate cardiac hypertrophy and heart failure requiring transplantation in middle age [50]. At early disease stage, R92W-TnT and R403Q-MyHC mouse hearts are characterized by distinct cardiac phenotypes and metabolic alterations impacting phospholipid metabolism which contrast the well-accepted notion of enhanced glycolysis in the setting of hypertrophic remodeling (**Figure 5**). We observed a notable shift in the total abundance and acyl-chain composition of cardiolipin (CL) in mutant TnT hearts. Incorporation of FA18:0 and FA18:1 in CL was increased, when compared to FA18:2, in TnT mutants. Cardiolipin is the most critical phospholipid of the inner mitochondrial membrane and is directly involved in mitochondrial function [51]. Together with PE, CL interacts and stabilizes respiratory chain complexes, ATP synthase, and supports mitochondrial cristae formation [52]. Studies on human and murine tissue demonstrate that CL acyl-chain composition is highly tissue specific [52–54]. Heart tissue is characterized by preferential incorporation of linoleic acid (FA18:2) [55]. Our R92W-TnT mutant model exhibits increased abundance of FA18:0 and FA18:1 in CL suggesting that the incorporation of FA18:2 is impaired. Alterations in the abundance and acyl-chain composition of CL in mitochondrial membranes can have functional consequences on electron transport chain function and ATP synthesis. The changes in mitochondrial phospholipid composition detected by our study in TnT mutant hearts could explain the molecular cardiac phenotype that we previously described [13] at early disease stage, characterized by lower mitochondrial complex I RCR, abnormalities in mitochondrial Ca^2+^handling and reduced mitochondrial DNA copy number [13].

Clinically, HCM patients are classified into 3 hemodynamic groups, namely, obstructive, non-obstructive and labile obstructive, based on the presence/absence of LV obstruction at rest and with provocation [43]. Patients can have non-obstructive hemodynamics at early disease stage that can either progress to obstructive HCM, or continue as non-obstructive HCM at established disease stage. Patients undergo septal myectomy [14, 56] when they have symptomatic LV obstruction that is unrelieved by medical therapy. Recent multi-omics profiling of myectomy tissue from HCM patients reported dysregulation of fatty acid oxidation, Krebs cycle, oxidative phosphorylation, and energy compromise [27, 28], which is similar to R92W-TnT hearts at early disease stage. Human HCM hearts showed a significant decrease in the abundance of acylcarnitines [27, 28], which is also evident in R92W-TnT and R403Q-MyHC mouse hearts. Additionally, human HCM myocardium demonstrated increases in PCs, PEs, lyso-PCs, lyso-PEs, ceramides, reduction in TAG, and increased lipid peroxidation [28], which is similar to our results in R92W-TnT mouse hearts. The most notable difference between our results in HCM mouse models and HCM patient studies, is the presence of genotype-specific metabolic remodeling in mouse hearts, and absence of correlation between genotype and cardiac metabolomic profile in human myectomy tissue [27–29]. There are several possible reasons for the differences in results between mouse and human metabolomics studies, including *1)* disease stage: early disease stage in HCM mouse models and established disease stage in HCM patients; *2)* cardiac physiology: non-obstructive HCM in mouse models, and obstructive HCM in patients undergoing septal reduction surgery (myectomy) [57]. These results lead us to hypothesize that LV obstruction induces a convergence to a common cardiac metabolic phenotype, in HCM.

### Limitations

Our studies characterizing cardiac metabolomics at early disease stage were only conducted in mouse models because it is unethical to obtain biopsies from asymptomatic patients with early stage HCM. Non-invasive investigation of cardiac metabolism that combine cardiac metabolic flux imaging with plasma profiling in early-stage human HCM could be helpful to confirm our results of genotype-specific metabolic remodeling, and to examine the association between metabolic, structural and functional remodeling in human HCM.

Untargeted metabolomics is inherently limited by the large number of unidentified metabolites. These compounds are frequently products of biotransformation and unspecific enzymatic reactions that physiologically occur but lead to ‘unknowns’ in LC-MS or GC-MS profiling scans. These compounds are not integrated in curated databases (e.g., HMDB, PubChem, KEGG) and often verified standards are missing. Furthermore, lipids are underrepresented in pathway-based databases which limits pathway enrichment analysis. Therefore, we expanded our analysis using CardioNet which includes cardiac-specific metabolic and lipidomic pathways.

Another limitation of this study is the exclusion of unknown metabolites in our analysis. More than 80% of metabolites are unknown in untargeted metabolomics workflows and structural resolution is limited. Therefore, we are reporting our entire dataset in the supplementary materials to engage with the scientific community and address this chemical complexity. Lastly, due to lack of method development targeted assays for nicotinamide adenine dinucleotide (NAD) such as NAD+/NADH could not be utilized in this study. Our metabolomics analysis yields high confidence data including MS and MS/MS data which provides a resource for the field.

## CONCLUSIONS

Multi-omics investigation, including comprehensive untargeted cardiac metabolomics and computer simulations revealed allele-specific differences in cardiac metabolism, that could underlie differences in the morphologic, functional and molecular cardiac phenotype at early disease stage. R92W-TnT hearts demonstrated reduction in energy provision and oxidative-stress induced phospholipid remodeling, whereas R403Q-MyHC hearts had preserved energy metabolism. Our results in mouse models demonstrate the need for in vivo imaging of cardiac metabolism in early-stage human HCM to enable individualized interventions that mitigate oxidative stress and deleterious phospholipid remodeling in HCM hearts.

## ACKNOWLEDGEMENTS

This work was supported by funding from the Department of Defense (PR171587 to MRA), West Coast Metabolomics Center Pilot Award (NIH U2C ES030158), National Institutes of Health (R00-HL-141702 to AK, R35GM145352 to RF), and the Leukemia Research Foundation (Grant#941997 to AK). The content is solely the responsibility of the authors and does not necessarily reflect the official views of the National Institute of Health. We are grateful to Dr. Gabriela V. Greenland for assistance with colony management, and Dr. Junaid Afzal for assistance with sample preparation.

## AUTHOR CONTRIBUTIONS

Conceptualization: MRA and AK. Methodology: MRA, OF, AV and AK. Experiments and Data analysis: AV, SF, YL, DAG, AK, TT and MRA. Writing and editing of manuscript: AK, AV, TT, OF, RF and MRA. Data curation: DAG, AV, and AK. Visualization: DAG, AK and AV. Computational flux analysis and CardioNet Modeling: AK. Funding acquisition: MRA and OF. Resources: OF, MRA, and AK.

## DATA AVAILABILITY

Raw files and unprocessed mass spectrometry data is available at the National Metabolomics Data Repository (Metabolomics WorkBench). Databased entry ID: ST002801 (GC-TOF), ST002815 (HILIC) and ST002817 (Lipidomics). The Sequence Read Archive (SRA) accession number for the mRNA-seq library reported here is SRP083078.

## METHODS

### Transgenic Mouse Models of HCM

Transgenic male C57BL/6 mice expressing the R403Q mutation in the α-Myosin Heavy Chain (MyHC) gene were kindly provided by Dr. Leslie Leinwand (University of Colorado Boulder). The R403Q-αMyHC mice were bred on a CBA/B16 (F1) cross background [32]. Transgenic male C56BL/6 mice bearing a c-myc-tagged murine TnT containing the R92W mutation were kindly provided by Dr. Jill Tardiff (University of Arizona). The R92W-TnT mouse is an F1 cross between FVB/N and C57/B6 strains [9, 58]. The R92W-TnT and R403Q-MyHC mice were backcrossed to C57BL/6 for >10 generations. Male mice were weaned and genotyped at the age of 4 weeks by PCR-amplified tail DNA. All studies were conducted at 5 weeks of age.

### Mouse phenotyping by echocardiographic imaging

Cardiac phenotyping was performed by echocardiography, using a Vevo 3100 platform and MX550D 40Mhz probe (VisualSonics, Toronto, Canada). Five-week-old R92W-TnT and R403Q-MyHC mice and littermate controls (WT) were anesthetized using inhaled isoflurane (3 v/v% for induction and 0.5-1% v/v% for maintenance). Parasternal long (PLSAX)- and short-axis (SAX) were recorded in B-mode and M-Mode. To evaluate diastolic function, pulsed-wave (PW) Doppler of mitral valve inflow, and left atrial area were recorded using the apical 4-chamber view. Heart rate was >450 bpm during recording of systolic parameters [59] and >400 bpm during assessment of diastolic function. Data was analyzed using the Vevo Image Lab Software (Version 5.6.0, VisualSonics, Toronto, Canada). Statistical Analysis was carried out using R (version 4.1.0) using the package gtsummary (version 1.4.1).

### Heart Sample Extraction for Metabolomics Analysis

#### Reagents

Water (H_2_O) (Optima LC/MS grade), methanol (MeOH) (Optima LC/MS grade), acetonitrile (ACN) (Optima LC/MS grade), 2-propanol (IPA) (Optima LC/MS grade), chloroform (CHCl_3_) (HPLC grade), hexane (Certified ACS), ammonium acetate (AA), methyl tert-butyl ether (MTBE), toluene (Tol) (HPLC grade 99.9%), formic acid (FA), ammonium formate (AmFo), Val-Tyr-Val, methoxyamine hydrochloride (MeOX), N-methyl-N-(trimethylsilyl)-trifluoroactamide (MSTFA), and pyridine (Anhydrous, 99.8%) were all purchased from Fisher Scientific (Hampton, NH, USA). Fatty acid methyl esters (FAMEs) are a mixture made from 13 standards that were purchased from both Fisher Scientific and Sigma-Aldrich. 12-(cyclohexylcarbamoylamino) dodecanoic acid (CUDA) was purchased from Cayman Chemical Company (Ann Arbor, MI, USA). Internal standards for the quality control (QC) mix for the lipidomics extraction were purchased from Nu-Check Prep, Inc. (Elysian, MN, USA), Avanti Polar Lipids (Alabaster, AL, USA), CDN Isotopes (Quebec, Canada), and Sigma-Aldrich. The internal standards in the resuspension solvent for HILIC analysis were purchased from CDN Isotopes, Cayman Chemicals (Ann Arbor, MI USA), Sigma-Aldrich, Toronto Research Chemicals, Inc. (Ontario, Canada), and Cambridge Isotope Laboratories, Inc. CIL Cambridge (Tewksbury, MA, USA).

#### Study design

Mice were euthanized by cervical dislocation, between 12 PM to 2 PM, following which the heart was rapidly excised, washed in ice cold PBS, flash frozen in liquid nitrogen, and stored at -80 °C until processing for metabolomics. Whole hearts were first homogenized in 100 μL ACN using 4 – 3 mm stainless steel grinding balls for 30 sec at a time for two rounds at 15,000 rotations per minute (rpm). Homogenized hearts were lyophilized overnight until a dry powder was obtained. Extraction for the lipidomics analysis was carried out using the Matyash et al. protocol [60]. Ice-cold (975 μL) of 3:10 (v/v) MeOH/MTBE with QC mix containing 19 internal standards was added to 5 mg of freeze-dried samples. Samples were mixed for 10 sec and then shaken for 5 min at 4°C; 188 μL of LC-MS grade H_2_O was added, mixed for 20 sec, and centrifuged for 2 min at 14,000 relative centrifugal force (rcf); 350 μL of the upper organic phase was separated into two new tubes; 100 μL from each sample analysis tube was transferred to a new tube to create a pooled sample. Samples were dried down and re-suspended with 110 μL of 9:1 (v/v) MeOH:Tol with 50 ng/mL CUDA before analysis; 50 μL was aliquoted into two separate amber glass vials with microvolume inserts for an injection in each mode. For HILIC analysis, 120 μL of the bottom aqueous phase from the lipidomics extraction mentioned above was split into two tubes. Samples were dried down and re-suspended in 100 μL of 80:20 (v/v) ACN:H_2_O with internal standards. The resuspension solvent for HILIC contains 42 internal standards. Samples were then mixed for 10 sec, sonicated for 5 min, and centrifuged for 2 min at 16,100 rcf. 90 μL was aliquoted into glass amber vials with microinserts prior to injection. For GC-TOF analysis, extraction and derivatization were carried out using the methods described in Fiehn et al [23–25]. One milliliter of degassed, -20°C pre-chilled extraction solvent of 3:3:2 (v/v/v) ACN:IPA:H_2_O was added to 4 mg of freeze dried sample. Samples were mixed for 10 sec, shaken for 5 min at 4°C, and then centrifuged for 2 min at 14,000 rcf. Two aliquots of 475 μL of the supernatant were separated into two tubes; 500 μL of 50:50 (v/v) ACN:H_2_O was added to dried down samples to clean up excess proteins then mixed for 10 sec and centrifuged for 2 min at 14,000 rcf; 475 μL of the supernatant was transferred to a new 1.5 mL tube which was then dried down to complete dryness. For derivatization, 10 μL MeOX solution was added to the sample and shaken for 1.5 hrs at 30°C. For trimethylsilylation, 91 μL of MSTFA with FAMEs internal standards mixture was added to each sample. Samples were centrifuged for 2 min at 14,000 rcf and then transferred to a GC crimp top glass vial with microvolume inserts and immediately capped with GC vial crimp caps for analysis.

### Instrumentation and metabolomic profiling

#### LC-MS/MS analysis: Lipidomics and HILIC analysis

Complex lipids were separated on a Waters Acquity UPLC CSH C18 column (100 × 2.1 mm; 1.7 μm) coupled to a Waters Acquity UPLC CSH C18 VanGuard pre-column (5 × 2.1 mm; 1.7 μm). The column was maintained at 65°C with a flow rate of 0.6 mL/min. The positive ionization mobile phases consisted of (A) 60:40 (v/v) ACN:H_2_O with 10 mM ammonium formate and 0.1% formic acid and (B) 90:10 (v/v) IPA/ACN with 10 mM ammonium formate and 0.1% formic acid. For negative mode, 10 mM ammonium acetate was used as the modifier. The following gradient was used: 0 min 15% B; 0-2min 30% B; 2-2.5 min 48% B; 2.5-11 min 82% B; 11-11.5 min 99% B; 11.5-12 min 99% B; 12-12.1 min 15% B; 12.1-15 min 15% B. 0.5 μL (ESI+) and 5.0 μL (ESI-) of sample was injected. The Agilent 6530 Q-TOF mass spectrometr (MS) coupled to an Agilent 1290 Infinity ultra-high-performance liquid chromatography (UHPLC) was operated in both positive and negative electrospray ionization (ESI) mode. Full MS1 data was acquired on pooled samples. The following parameters for Full MS1 were used mass range *m/z* 120 – 1700 (ESI+) and *m/z* 60 – 1700 (ESI-), gas temperature 325°C, drying gas (nitrogen) flow rate 8 L/min, nebulizer 35 psig, sheath gas (nitrogen) flow rate 11 L/min, sheath gas temperature 350°C, VCap 3500 V, and nozzle voltage 1000 V. AutoMS/MS is acquired using the following parameters: mass ranges using Static Exclusion Range list *m/z* 120 – 700, *m/z* 700 – 800, *m/z* 800 – 880, and *m/z* 880 – 1700 (ESI+) and *m/z* 60 – 700, *m/z* 700 – 800, *m/z* 800 – 880, and *m/z* 880 – 1700 (ESI-). *Use Formula* option was used to ramp collision energies (CE) from low to high, the following parameters were used: slope 3 (ESI+) and 4 (ESI-), Offset 2.5 (ESI+/-), and charge All (ESI +/-). Data were acquired in centroid.

Biogenic amines were analyzed by HILIC LC-MS/MS analysis. Metabolites were separated on a Waters Acquity UPLC BEH Amide column (150 × 2.1 mm; 1.7 μm) coupled to an Acquity UPLC BEH Amide VanGuard pre-column (5 × 2.1 mm; 1.7 μm). The column was maintained at 40°C with a flow rate of 0.4 mL/min. The mobile phase consisted of (A) H_2_O with 10 mM ammonium formate and 0.125 % formic acid at pH 3 and (B) 95:5 (v/v) ACN/H_2_O with 10 mM ammonium formate and 0.125% formic acid at pH3. The following gradient was used: 0 min 100% B; 0-2 min 100% B; 2-7 min 70% B; 7-7.9 min 40% B; 9.5-10.25 min 30% B; 10.25-12.75 min 100% B; 16.75 min 100% B. 1.0 μL (ESI+) of sample was injected. The SCIEX 6600 TripleTOF MS coupled to an Agilent 1290 Infinity UHPLC was operated in positive ESI mode using the following parameters: mass range 50 – 1500 Da (MS1) and 40 – 1000 Da (MS/MS using Information Acquistion), CE 35 V, Duration 13.999 min, Cycle Time 0.5002 sec, and Accumulation Time 0.250017 sec. The following source parameters with arbitrary units were used: curtain gas (CUR) 35, IonSpray Voltage (IS) 4000, temperature (TEM) 500, Ion Source Gas (GSI) 50, Ion Source Gas 2 (GS2). Data was acquired in centroid.

#### GC-MS analysis: Primary metabolism by GC-TOF

Data for primary metabolites was acquired on the LECO Pegasus High-Throughput TOF-MS coupled to an Agilent 7890A gas chromatograph with Gerstel MultiPurpose Autosampler. Separation was achieved on Restek RTX-5Sil MS column (30 m length, 0.25 mm i.d., and 0.25 μm 95% dimethyl 5% diphenyl polysiloxane film) with a 10Lm guard column. The injection volume was 0.5 μL at 250L°C. The GC used the flowing oven program held for 1 min at 50 °C, ramped to 330 °C at 20 °C/min, and held for 5 min before cool-down. The transfer line temperature was 280 °C. Spectra were recorded with the following parameters: mass range 85 – 500 u, electron ionization (EI) mode 70 eV, filament temperature 250 °C, acquisition time 20 min, and scan rate 17 spectra/sec.

### Data processing

#### LC-MS/MS

Untargeted LC-MS/MS data was analyzed by MS-DIAL software version 3.52 for lipidomics and 3.06 for HILIC [26]. Detailed parameter settings for lipidomics and HILIC data processing are listed in **Supplemental Table 1**. Metabolite annotations were done using our in-house developed mass-to-charge-retention time (m/z-RT) libraries. MS/MS spectral matching was performed using freely available MS/MS libraries obtained from Mass Bank of North America (MoNA) (www.massbank.us), and commercially available NIST17 MS/MS library.

#### GC-MS

The peak and compound detection or deconvolution were performed with the LECO ChromaTOF software version 4.50.1. Spectra were matched against the FiehnLib mass spectral and retention index library [39]. Post-curation and peak replacements were performed with the in-house developed BinBase software [61] and the sample matrix with all known and unknown compounds exported to a Microsoft EXCEL sheet. Peak heights were normalized by mTIC normalization, mTIC is defined as the sum of all the peak heights for all identified metabolites.

### Mathematical modeling of HCM heart metabolism

*In silico* simulations were performed using the metabolic network of the cardiomyocyte CardioNet [30, 48, 62]. Mathematical modelling has previously been used to study the dynamics of cardiac metabolism in response to stress, [30, 63, 64] and CardioNet has been successfully applied to identify limiting metabolic processes and estimate flux distributions [62, 64]. Flux balance analysis (FBA) allows to estimate flux rates in a cellular model based on metabolic constraints that are defined by the extracellular environment (e.g. oxygen and nutrient supply), cellular demands (e.g. proliferation, contraction) and tissue type. This modeling approach combines biochemical network models with optimality problems, which describe different cost or benefit functions and allow us to include experimental data, for example metabolite levels, enzyme levels or flux rates. The advantage of flux balance analysis is that it considers system-wide effects of processes and allows us to assess metabolic limitations in an unbiased approach. We applied flux balance analysis to identify which reactions are involved in myocardial metabolic adaptations to R403Q-MyHC or R92W-TnT variants. Metabolic flux distributions were calculated using constrained based modeling. To calculate flux rate changes (*v_i_*), we constrained the model for each metabolite using experimentally determined metabolite concentrations (^1^H NMR) to maximize cardiac work reflected by ATP hydrolysis (v_ATPase_). Simulations were run with boundary conditions reflecting the circulating metabolite composition based on previously reported values and common metabolites present in plasma^65-67,46^. Based on these constrains we first determined flux distributions (*v_m_*) under sham operation (control) conditions. We then calculated fold-changes (FC) for experimentally-measured metabolite concentrations between control and R92W-TnT or R403Q-MyHC, and used these fold-changes to further constrain fluxes (*v_m_*) for the synthesis and/or degradation of intracellular metabolites.

We included fold-changes (FC) based on the assumption that changes in metabolite concentrations under experimental conditions are accompanied by a proportional increase or decrease in the respective flux for the metabolite pool. By using metabolite level changes (fold changes) to estimate flux rate changes (v_FC_), we imply that the altered steady-state concentrations of metabolites are reflected in the newly evolved flux state and potentially limit metabolic functions. The following flux balance analysis was applied to identify steady-state flux distributions that agree with applied substrate uptake and release rates, and changes in metabolite pools:

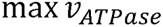

subject to

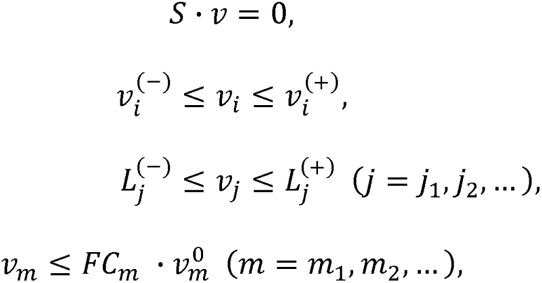

where *v_i_* denotes the flux rate change through reaction *i*, *v_j_* denotes the measured uptake or secretion rate through reaction *j*, *S* is the stoichiometric matrix, and 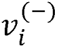 and 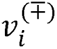are flux constraints. The GUROBI LP solver was used to find the solution to the FBA problems.^68^ The logarithm of the metabolic flux rate values is presented in the form of heatmaps. Metabolic reactions are clustered according to their association to metabolic pathways and plot colours indicate estimated flux rates for each metabolic reaction. All reactions and their metabolic subsystems, classified in the Kyoto Encyclopedia of Genes and Genomes database [69].

### Statistical Analysis

Mann-Whitney U test was used on each metabolite to test the significance of two pathogenic variants in sarcomeric protein genes that lead to HCM: mutant R92W-TnT hearts (TnT) versus littermate control TnT mice (TnT vs. WT) and R403Q-MyHC mice (MyHC) versus littermate control mice (MyHC vs. WT). The fold change of each comparison was calculated as the group median ratio. To control the false discovery rate (FDR), we adopted the Benjamini-Hochberg FDR correction procedure [70]. We performed chemical enrichment analysis (ChemRICH) [40] using the p-values and fold changes to access chemical classes significantly altered (p<0.05) in each comparison. ChemRICH provides enrichment analysis based upon chemical structure and not pre-defined pathways which can be inherently flawed and does not rely upon background databases for statistical calculations. Metabolite cluster significance was calculated using Kolmogorov–Smirnov test. Metabolites were considered significant using the Mann-Whitney U test if p< 0.05. Downstream analysis of processed metabolomics data relied on (1) the use of non-parametric statistical methods, which do not make assumptions on the underlying distribution of the data, and (2) the use of littermate control mouse heart tissue as a “control” or “reference” metabolome by which to identify specific metabolic changes due pathogenic variants in sarcomeric protein genes. We annotated metabolites to identifiers of several key databases including KEGG, HMDB, PubChem IDs and CardioNet [30].

---

**Supplemental Fig S1.**
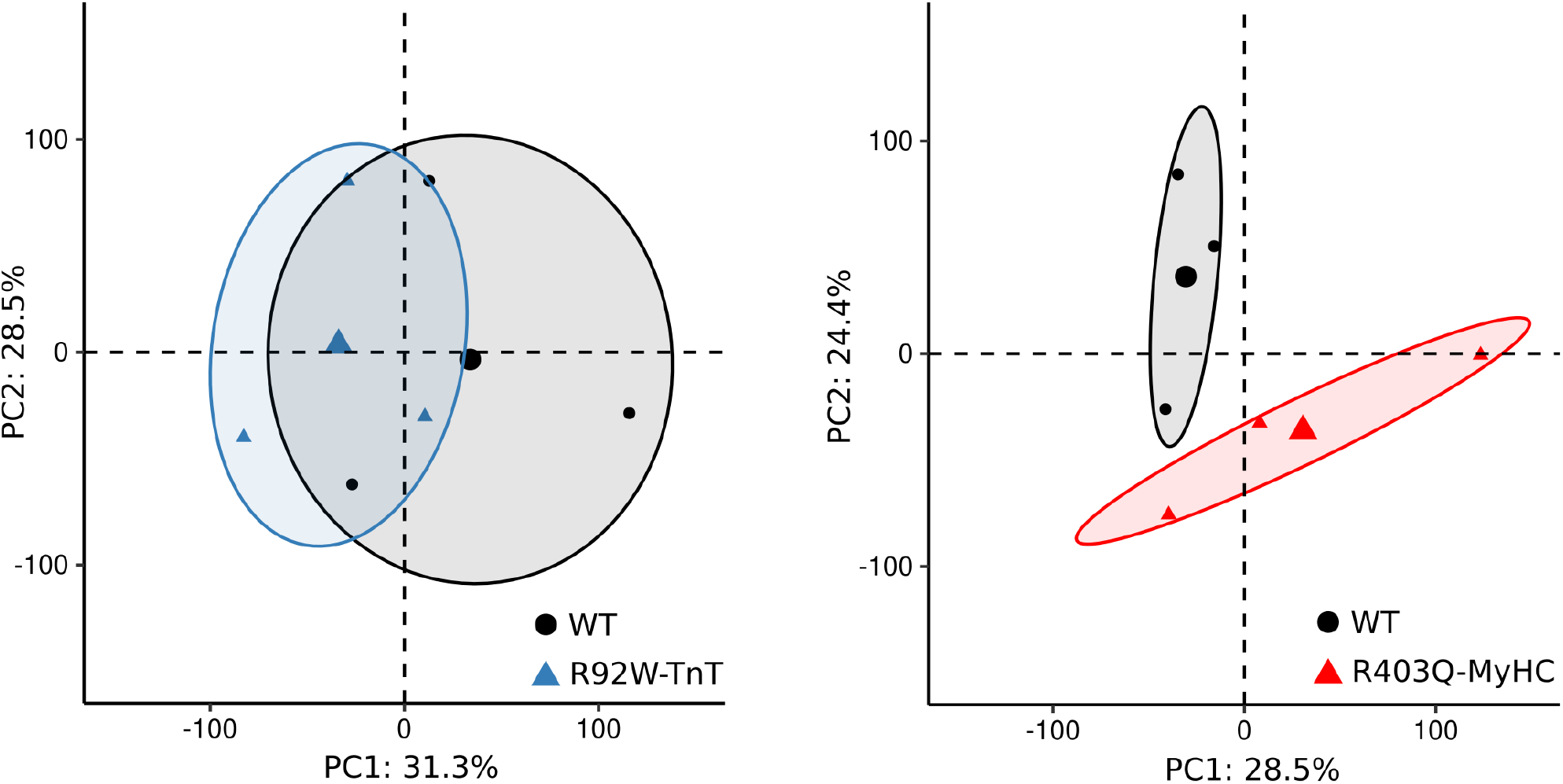
PCA plot of gene expression data in R92W-TnT and R403Q-MyHC hearts compared to littermate control hearts (WT) at early disease stage. Dimensions 1 and 2 explain >50% of variance in both mutant hearts at 5 weeks of age.

**Supplemental Figure S2.**
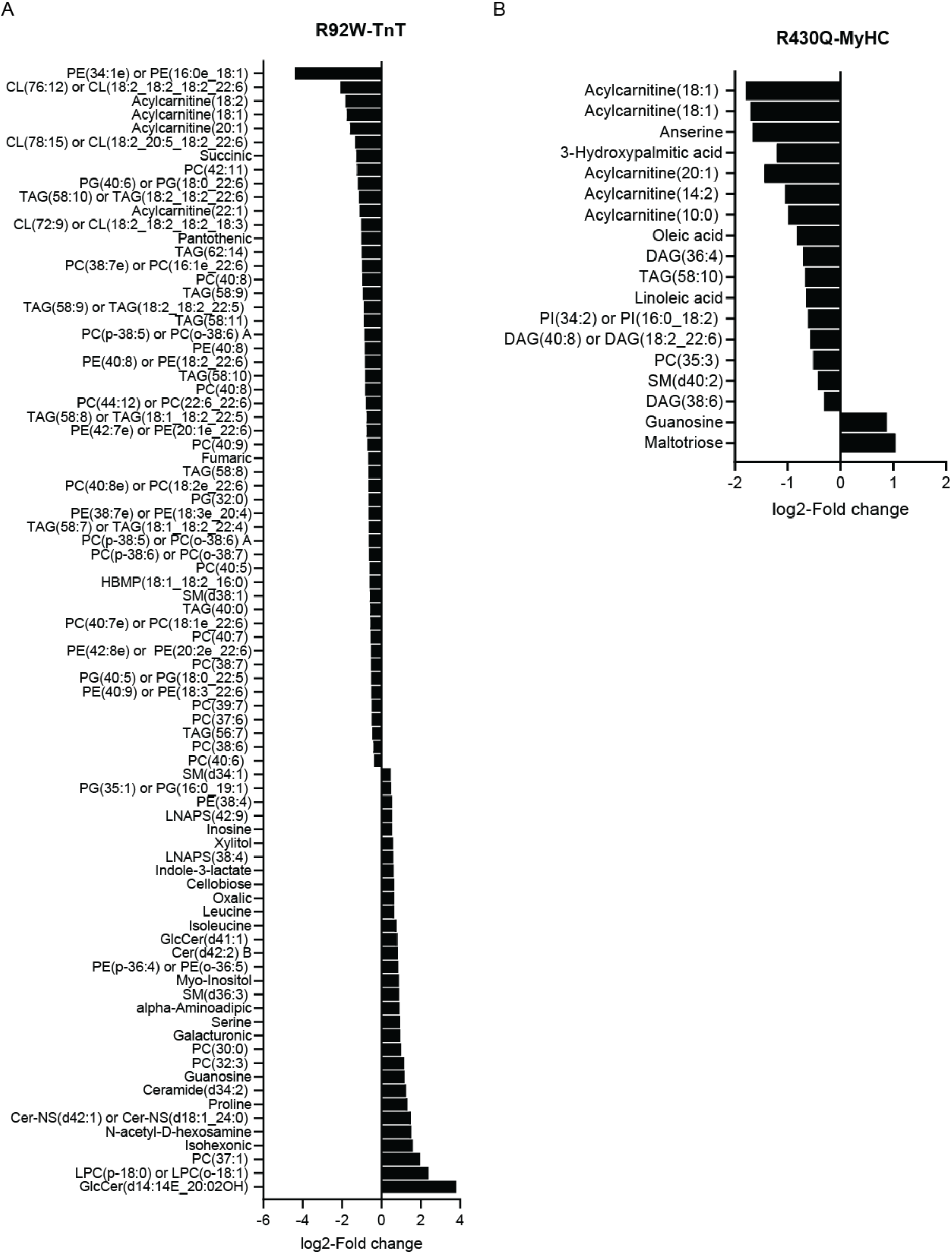
Analysis of metabolite and lipid distributions in R92W-TnT and R403Q-MyHC mouse hearts at early disease stage. We find major alterations in (**A**) R92W-TnT hearts for phospholipids, including cardiolipin (CL), ceramide (Cer), lyso-phosphatidylcholine (LPC), phosphatidyl-choline (PC), -ethanolamine (PE), and –glycerol (PG) species. For visualization purposes, only significantly altered metabolites and lipids with an adjusted p-value<0.05 and a log2-fold change (>2 or <-0.5) are depicted. (**B**) R403Q-MyHC hearts show significant alterations in acylcarnitine species and total abundance of NAD (p-value<0.05; log2- Fold change >2 or <-0.5).

**Supplemental Figure S3.**
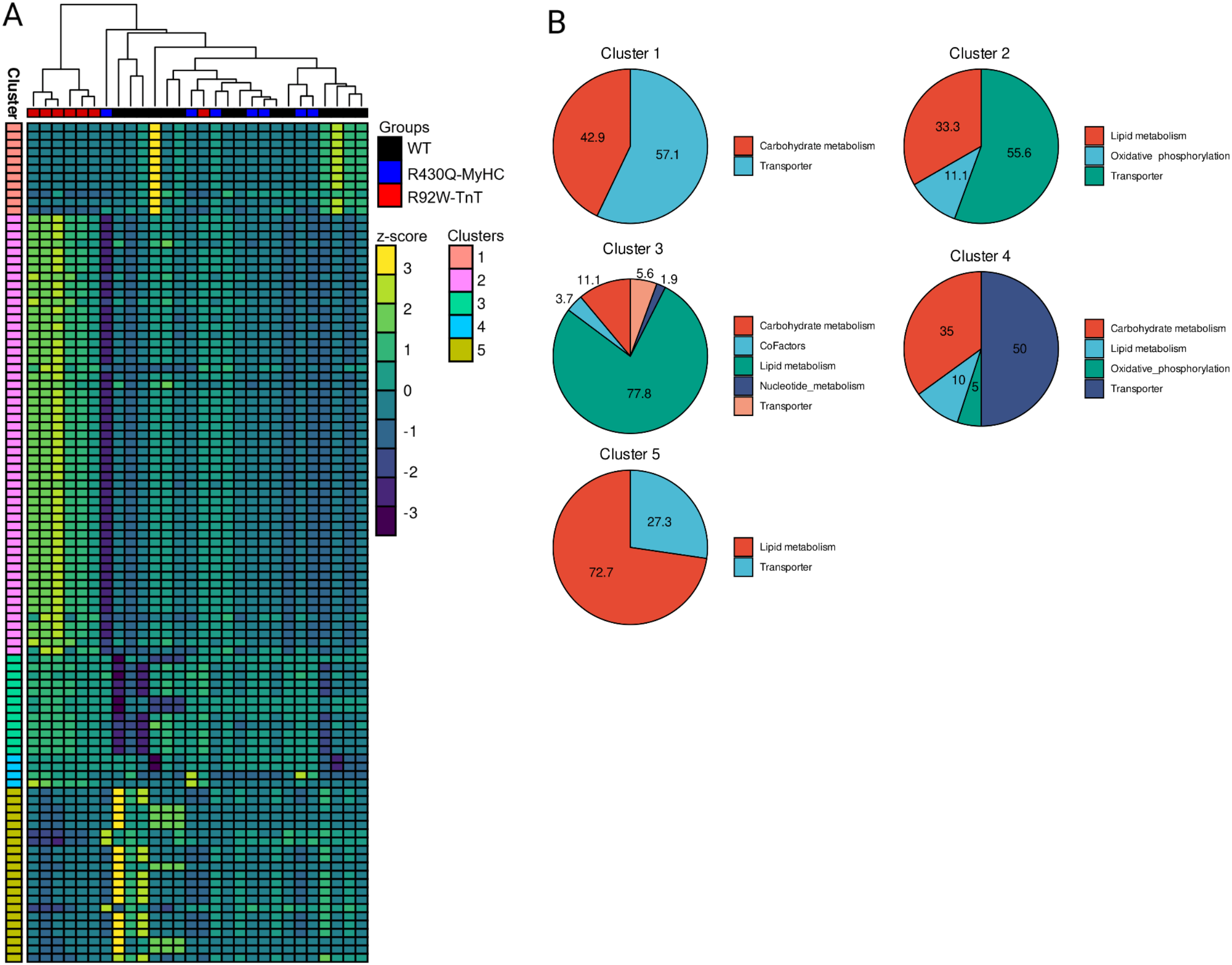
Mathematical modelling of R92W-TnT and R403Q-MyHC mouse hearts at early disease stage. (**A**) Unsupervised hierarchical cluster analysis of calculated metabolic flux rates. Data was normalized using z-scores. (**B**) Pathway distribution across identified cluster related to panel A.

