## Supplemental File for "Lipid metabolism drives allele-specific early-stage hypertrophic cardiomyopathy"

**Supplementary Data**

**Allele-specific differences in cardiac metabolism in early-stage hypertrophic cardiomyopathy: Metabolomics results from two mouse models at early disease stage**

Arpana Vaniya^1*^, Anja Karlstaedt^2,3*^, Damla Ates Gulkok^4^, Tilo Thottakara^4^, Yamin Liu^4^, Sili Fan^1^, Hannah Eades^4^, Ryuya Fukunaga^5^, Hilary J. Vernon^6, 7^, Oliver Fiehn^1^^, M. Roselle Abraham^4^^

^1^West Coast Metabolomics Center, University of California, Davis, Davis, CA

^2^Department of Cardiology, Smidt Heart Institute, Cedars-Sinai Medical Center, Los Angeles, CA

^3^Department of Biomedical Sciences, Cedars-Sinai Medical Center, Los Angeles, CA

^4^Hypertrophic Cardiomyopathy Center of Excellence, Division of Cardiology, University of California San Francisco, San Francisco, CA

^5^Department of Biologic Chemistry, Johns Hopkins University, Baltimore, MD

^6^McKusick Nathans Department of Genetic Medicine, Johns Hopkins University, Baltimore, MD

^7^Department of Pediatrics, Johns Hopkins University, Baltimore, MD

*Contributed Equally

Arpana Vaniya, PhD

Anja Karlstaedt, MD PhD

Address correspondence to:

^^^M. Roselle Abraham, M.D.

^^^Oliver Fiehn, PhD

Keywords: Untargeted metabolomics, lipidomics, hypertrophic cardiomyopathy, RNAseq, HCM mouse models

Table S1. Parameter settings for LC-MS/MS data processing

| Parameters | Value (POS Mode) | Value (NEG Mode) | Value (POS Mode) |
| --- | --- | --- | --- |
| Platform | Lipidomics | Lipidomics | HILIC |
| MS-DIAL Version | 3.52 | 3.52 | 3.06 |
| Data collection parameters |  |  |  |
| Retention time begin | 0.5 | 0.5 | 0.5 |
| Retention time end | 12.5 | 12.5 | 14 |
| Mass range begin | 50 | 50 | 50 |
| Mass range end | 1700 | 1700 | 1700 |
| Centroid parameters |  |  |  |
| MS1 tolerance | 0.01 | 0.01 | 0.01 |
| MS2 tolerance | 0.025 | 0.025 | 0.025 |
| Isotope recognition |  |  |  |
| Maximum charged number | 2 | 2 | 2 |
| Data processing |  |  |  |
| Number of threads | 8 | 8 | 8 |
| Peak detection parameters |  |  |  |
| Smoothing method | LinearWeightedMovingAverage | LinearWeightedMovingAverage | LinearWeightedMovingAverage |
| Smoothing level | 3 | 3 | 3 |
| Minimum peak width | 5 | 5 | 5 |
| Minimum peak height | 1000 | 1000 | 1000 |
| Peak spotting parameters |  |  |  |
| Mass slice width | 0.1 | 0.1 | 0.1 |
| Exclusion mass list (mass & tolerance) | None | None | None |
| Deconvolution parameters |  |  |  |
| Sigma window value | 0.1 | 0.1 | 0.1 |
| MS2Dec amplitude cut off | 0 | 0 | 0 |
| Exclude after precursor | TRUE | TRUE | TRUE |
| Keep isotope until | 0.5 | 0.5 | 0.5 |
| Keep original precursor isotpes | FALSE | FALSE | FALSE |
| Exclude after precursor | TRUE | TRUE | TRUE |
| MSP file and MS/MS identification setting |  |  |  |
| MSP file | LipidMsmsBinaryDB-VS47-FiehnO.lbm2 | LipidMsmsBinaryDB-VS47-FiehnO.lbm2 | MoNA+NIST17.msp |
| Retention time tolerance | 100 | 100 | 100 |
| Accurate mass tolerance (MS1) | 0.01 | 0.01 | 0.01 |
| Accurate mass tolerance (MS2) | 0.05 | 0.05 | 0.05 |
| Identification score cut off | 80 | 80 | 80 |
| Using retention time for scoring | TRUE | TRUE | TRUE |
| Text file and post identification (retention time and accurate mass based) setting |  |  |  |
| Text file | CORE_posCSH_mzrt_041719.txt | CORE_negCSH_mzrt_041719.txt | TTOF_posHILIC_mzrt_091219.txt |
| Retention time tolerance | 0.1 | 0.1 | 0.1 |
| Accurate mass tolerance | 0.01 | 0.01 | 0.01 |
| Identification score cut off | 85 | 85 | 85 |
| Advanced setting for identification |  |  |  |
| Relative abundance cut off | 0 | 0 | 0 |
| Top candidate report | TRUE | TRUE | TRUE |
| Adduct ion setting | [M+H]+ | [M-H]- | [M+H]+ |
|  | [M+NH4]+ | [M+Cl]- | [M+NH4]+ |
|  | [M+Na]+ | [M+Hac-H]- | [M+Na]+ |
|  | [M+K]+ | [2M-H]- | [M+K]+ |
|  | [M+H-H2O]+ |  | [M+H-H2O]+ |
|  | [2M+H]+ |  | [2M+H]+ |
| Alignment parameters setting |  |  |  |
| Reference file | First sample | First sample | First sample |
| Retention time tolerance | 0.05 | 0.05 | 0.05 |
| MS1 tolerance | 0.015 | 0.015 | 0.015 |
| Retention time factor | 0.5 | 0.5 | 0.5 |
| MS1 factor | 0.5 | 0.5 | 0.5 |
| Peak count filter | 0 | 0 | 0 |
| QC at least filter | FALSE | FALSE | FALSE |
| Remove feature based on peak height fold-change | FALSE | FALSE | FALSE |
| Sample max / blank average | 5 | 5 | 5 |
| Sample average / blank average | 5 | 5 | 5 |
| Keep identified and annotated metabolites | TRUE | TRUE | TRUE |
| Keep removable features and assign the tag for checking | TRUE | TRUE | TRUE |
| Replace true zero values with 1/2 of minimum peak height over all samples | FALSE | FALSE | FALSE |
| Tracking of isotope labels |  |  |  |
| Tracking of isotopic labels | FALSE | FALSE | FALSE |
| Load Together with Alignment | Check | Check | Check |

Table S2. Compound classes using LC-MS/MS Lipidomics

| Class | Number of Metabolites | Percentage |
| --- | --- | --- |
| Unsaturated triglycerides | 125 | 22.40% |
| Unsaturated phosphatidylcholines | 75 | 13.44% |
| Ether phosphatidylethanolamines | 42 | 7.53% |
| Unsaturated phosphatidylethanolamines | 38 | 6.81% |
| Ether phosphatidylcholines | 32 | 5.73% |
| Cardiolipins | 25 | 4.48% |
| Unsaturated phosphatidylinositols | 25 | 4.48% |
| Unsaturated sphingomyelins | 24 | 4.30% |
| Unsaturated ceramides | 19 | 3.41% |
| Unsaturated phosphatidylglycerols | 19 | 3.41% |
| Unsaturated fatty acyls | 17 | 3.05% |
| Acylcarnitine | 14 | 2.51% |
| Diacylglycerols | 14 | 2.51% |
| Saturated fatty acyls | 12 | 2.15% |
| Saturated triglycerides | 11 | 1.97% |
| Unsaturated lysophosphatidylcholines | 9 | 1.61% |
| Saturated phosphatidylcholines | 7 | 1.25% |
| Galactosylceramides | 6 | 1.08% |
| N-acyl-lysophosphatidylethanolamine | 6 | 1.08% |
| Saturated lysophosphatidylcholines | 6 | 1.08% |
| Saturated sphingomyelins | 5 | 0.90% |
| N-acyl-lysophosphatidylserine | 4 | 0.72% |
| Cholesterol esters | 3 | 0.54% |
| Saturated ceramides | 3 | 0.54% |
| Unsaturated lysophosphatidylethanolamines | 3 | 0.54% |
| Unsaturated phosphatidylserines | 3 | 0.54% |
| Cholestane steroids | 2 | 0.36% |
| Ether lysophosphatidylcholines | 2 | 0.36% |
| Ganglioside GM3 | 2 | 0.36% |
| Saturated lysophosphatidylethanolamines | 2 | 0.36% |
| Hemibismonoacylglycerophosphate | 1 | 0.18% |
| Saturated phosphatidylethanolamines | 1 | 0.18% |
| Saturated phosphatidylglycerols | 1 | 0.18% |
| Total number of metabolites | 558 |  |

Table S3. Compound classes using GC-TOF and HILIC

| Class | Number of Metabolites | Percentage |
| --- | --- | --- |
| Amino acids and derivatives | 63 | 19.21% |
| Fatty Acyls | 44 | 13.41% |
| Carbohydrates and carbohydrate conjugates | 43 | 13.11% |
| Peptides | 16 | 4.88% |
| Imidazopyrimidines | 10 | 3.05% |
| Glycerolipids | 9 | 2.74% |
| Pyrimidine nucleotides | 8 | 2.44% |
| Purine nucleotides | 8 | 2.44% |
| Purine nucleosides | 8 | 2.44% |
| Benzene and substituted derivatives | 8 | 2.44% |
| Amines | 7 | 2.13% |
| Hydroxy acids and derivatives | 7 | 2.13% |
| Steroids and steroid derivatives | 7 | 2.13% |
| Pyrimidine nucleosides | 5 | 1.52% |
| Diazines | 5 | 1.52% |
| Indoles and derivatives | 5 | 1.52% |
| Dicarboxylic acids and derivatives | 5 | 1.52% |
| Pyridines and derivatives | 5 | 1.52% |
| Quaternary ammonium salts | 5 | 1.52% |
| Secondary alcohols | 4 | 1.22% |
| 5'-deoxy-5'-thionucleosides | 4 | 1.22% |
| Lactones | 3 | 0.91% |
| Organic carbonic acids and derivatives | 3 | 0.91% |
| Prenol lipids | 3 | 0.91% |
| Phenols | 3 | 0.91% |
| Phosphate esters | 3 | 0.91% |
| Peptidomimetics | 3 | 0.91% |
| Carbonyl compounds | 2 | 0.61% |
| Keto acids and derivatives | 2 | 0.61% |
| Non-metal oxoanionic compounds | 2 | 0.61% |
| Oxepanes | 2 | 0.61% |
| Cyclic alcohols and derivatives | 2 | 0.61% |
| Azoles | 2 | 0.61% |
| Flavin nucleotides | 2 | 0.61% |
| (5'->5')-dinucleotides | 2 | 0.61% |
| Sphingolipids | 2 | 0.61% |
| Organosulfonic acids | 1 | 0.30% |
| Nucleoside and nucleotide analogues | 1 | 0.30% |
| Phenylpropanoic acids | 1 | 0.30% |
| Glycerophospholipids | 1 | 0.30% |
| Tricarboxylic acids and derivatives | 1 | 0.30% |
| Butenolides | 1 | 0.30% |
| Tetrahydrofurans | 1 | 0.30% |
| Benzoxazines | 1 | 0.30% |
| Ribonucleoside 3'-phosphates | 1 | 0.30% |
| Carboximidic acids | 1 | 0.30% |
| Pyrroles | 1 | 0.30% |
| Porphyrins | 1 | 0.30% |
| Pteridines and derivatives | 1 | 0.30% |
| Benzothiazines | 1 | 0.30% |
| Alkaloids and derivatives | 1 | 0.30% |
| Trialkyl amine oxides | 1 | 0.30% |
| Total number of metabolites | 328 |  |

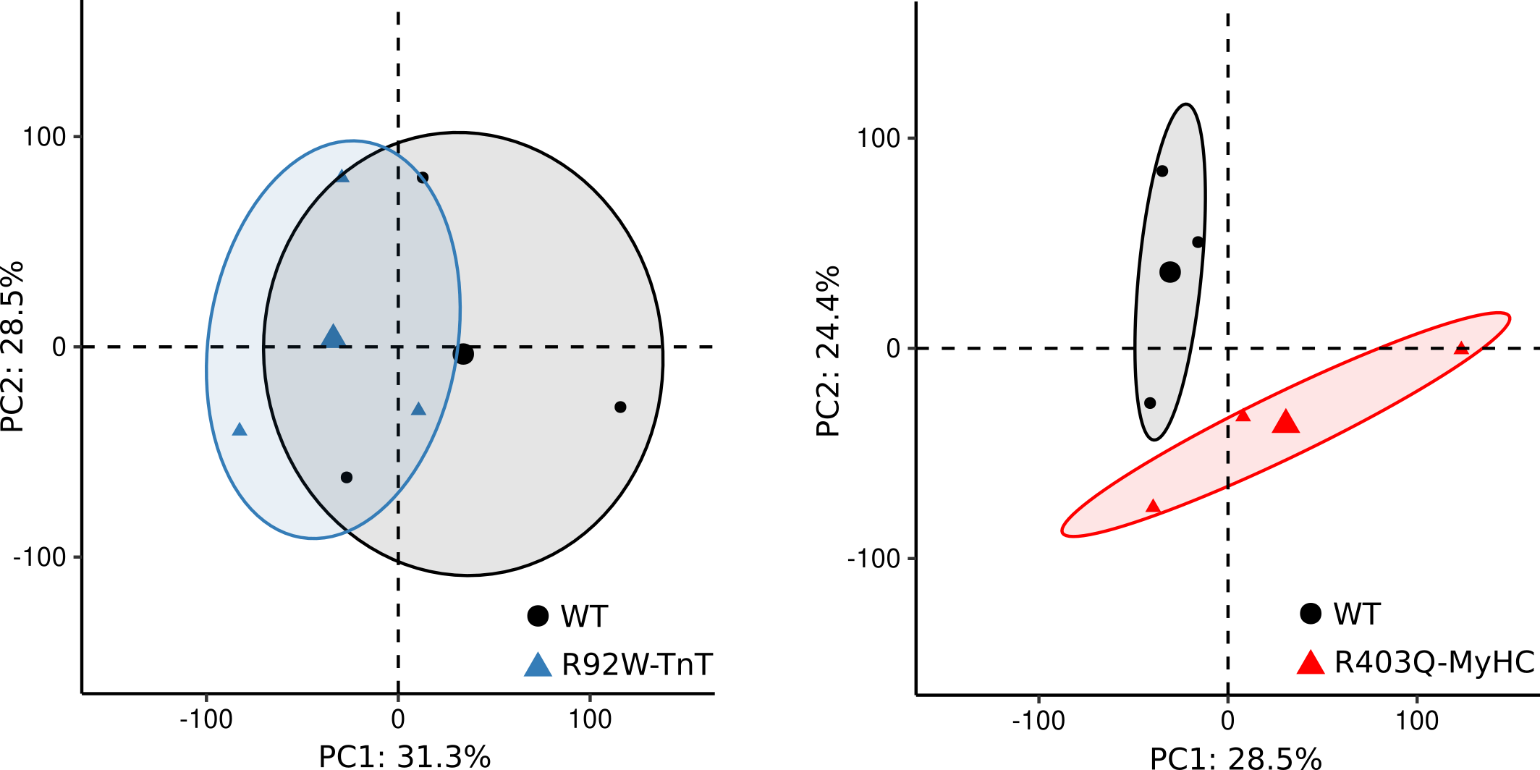

**Supplemental Figure S1. PCA plot for transcriptomics data in R92W-TnT and R403Q-MyHC mouse hearts at early disease stage.** Dimensions 1 and 2 explain >50% of the variance in both mutant hearts at 5 weeks of age.

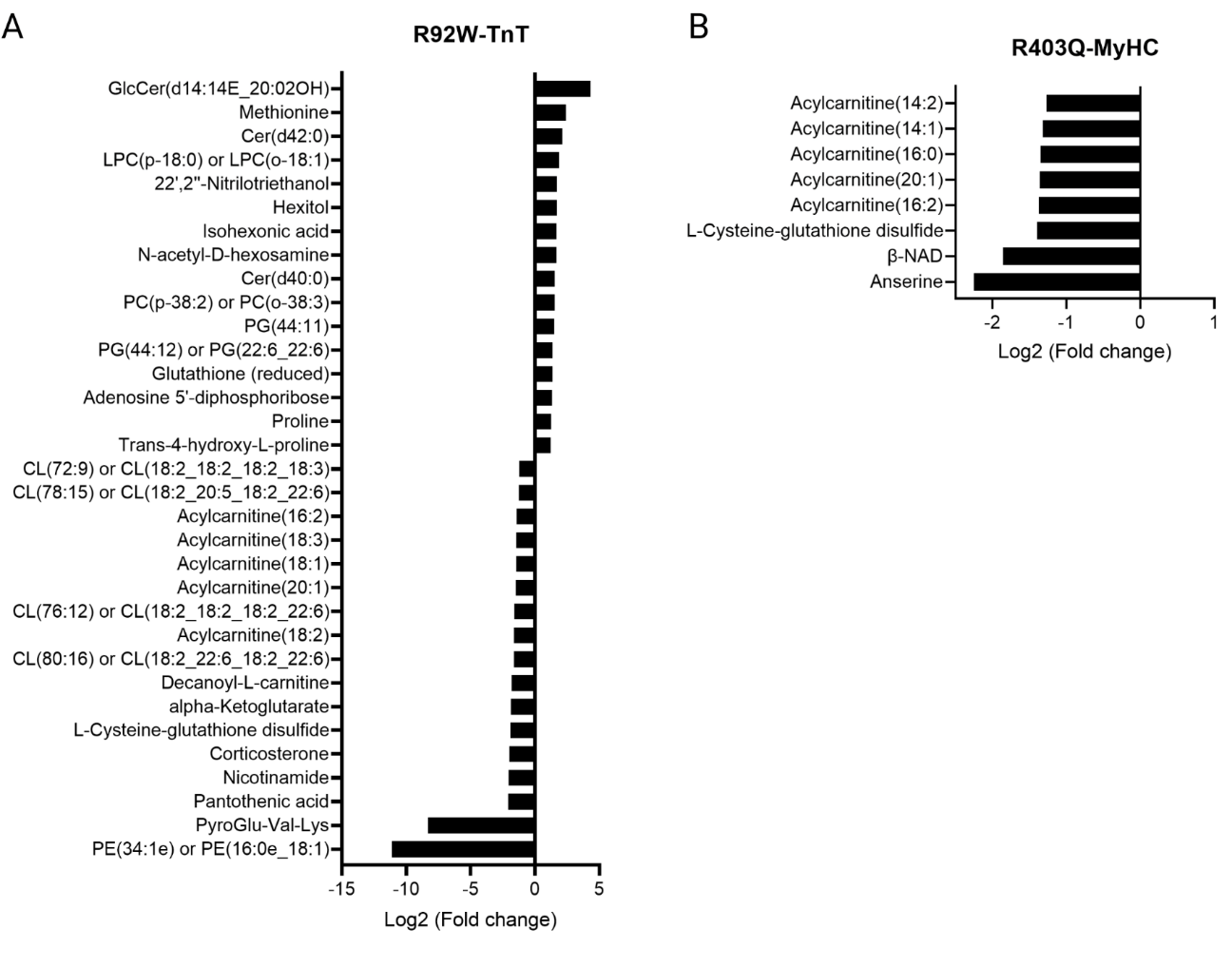

**Supplemental Figure S2. Analysis of metabolite and lipid distributions in R92W-TnT and R403Q-MyHC mouse hearts at early disease stage.** Significantly altered metabolites and lipids (p-value<0.05) are depicted as a function of log2-Fold change (>2 or <-0.5) compared to littermate controls (WT). We find major alterations in (**A**) R92W-TnT hearts for phospholipids, including cardiolipin (CL), ceramide (Cer), lyso-phosphatidylcholine (LPC), phosphatidyl-choline (PC), -ethanolamine (PE), –glycerol (PG) species. (**B**) R403Q-MyHC hearts show significant alterations in acylcarnitine species and total abundance of NAD.

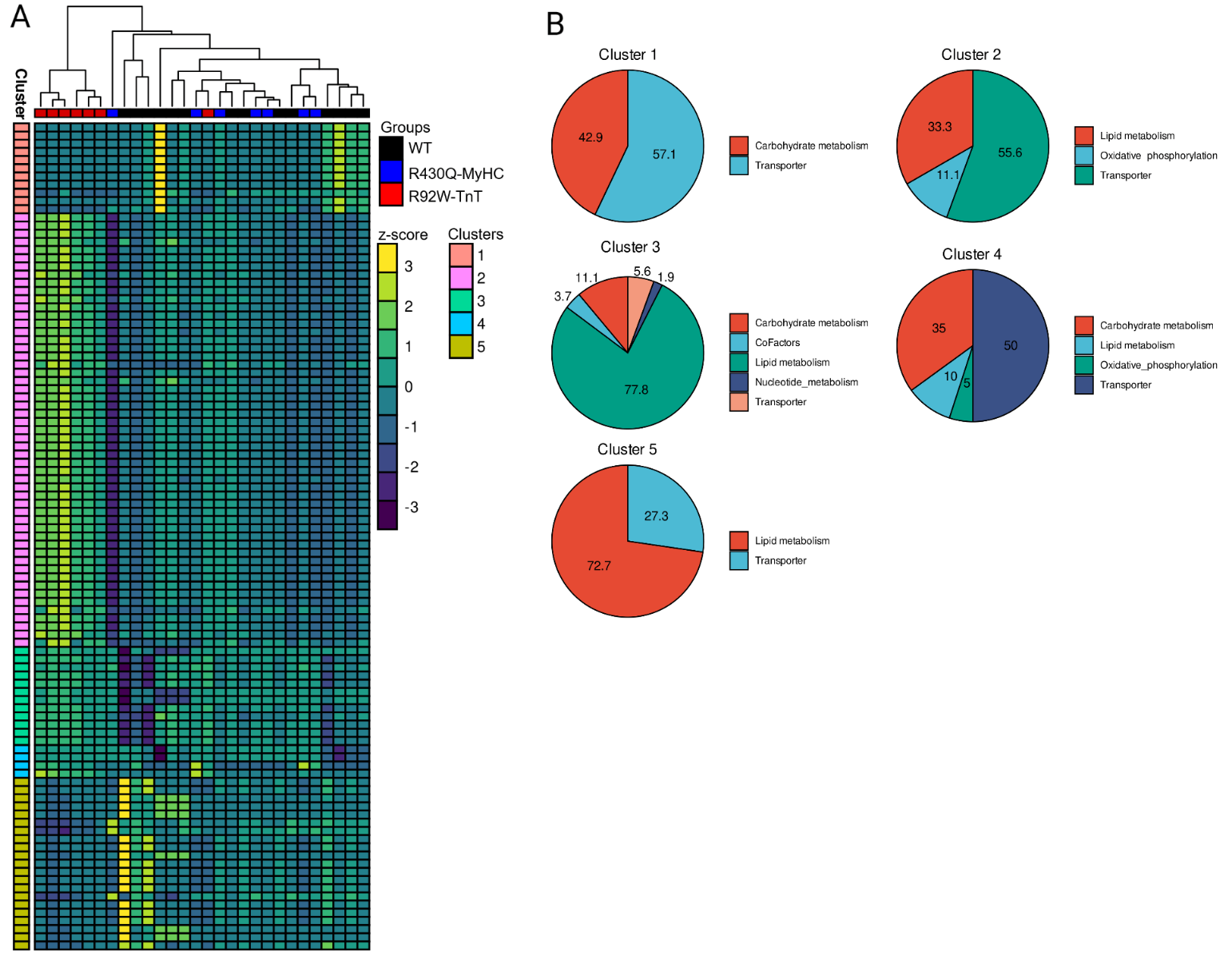

**Supplemental Figure S3.** Mathematical modelling of metabolism in R92W-TnT and R403Q-MyHC mouse hearts at early disease stage. (A) Unsupervised hierarchical cluster analysis of calculated metabolic flux rates. Data was normalized using z-scores. (B) Pathway distribution across identified cluster related to panel A.
